# Sphingolipids protect ergosterol in the *Leishmania major* membrane from sterol-specific toxins

**DOI:** 10.1101/2022.06.15.496300

**Authors:** Chaitanya S. Haram, Samrat Moitra, Rilee Keane, F. Matthew Kuhlmann, Cheryl Frankfater, Fong-Fu Hsu, Stephen M. Beverley, Kai Zhang, Peter A. Keyel

## Abstract

Susceptibility of *Leishmania* to the first line treatment amphotericin B remains poorly understood. Amphotericin B targets ergosterol, so one approach to improving drug efficacy and reducing side effects could be improving access to ergosterol. While the surface exposure of ergosterol in *Leishmania* is unknown, sterols in mammalian cells can be sheltered from sterol-binding agents by membrane components, including sphingolipids. Here, we tested the ability of the *Leishmania major* sphingolipids inositol phosphorylceramide (IPC), and ceramide to shelter ergosterol by preventing binding and cytotoxicity of the sterol-specific toxins streptolysin O and perfringolysin O using flow cytometry. In contrast to mammalian systems, *Leishmania* sphingolipids did not preclude toxin binding to sterols in the membrane. However, IPC interfered with cytotoxicity. Ceramide reduced perfringolysin O, but not streptolysin O, cytotoxicity in cells. Ceramide sensing was controlled by the toxin L3 loop. Ceramide was sufficient to protect *L. major* promastigotes from amphotericin B. We propose a mechanism whereby pore-forming toxins engage additional lipids like ceramide to determine the optimal environment to sustain pore formation. Thus, *L*. *major* offers a genetically tractable model organism for understanding toxin-membrane interactions. Furthermore, our findings suggest targeting ceramide may enhance the efficacy of ergosterol-targeting anti-leishmanial drugs.

**Abstract Importance:** Leishmaniasis is a neglected tropical disease with ∼1.5-2 million new cases and ∼70,000 deaths annually. One first-line treatment for leishmaniasis is liposomal amphotericin B, which is expensive and damages the kidneys. Cost and side effects can be minimized by improving efficacy. To improve efficacy, we must learn how amphotericin’s target—ergosterol—is protected by other components of *Leishmania*. The human ergosterol equivalent is protected by components called sphingolipids. We tested the ability of sphingolipids to protect ergosterol using pore-forming toxins. Pore-forming toxins use ergosterol to bind and kill *Leishmania*. Unlike human cells, toxins bound to ergosterol—indicating that they had access—when sphingolipids were present. However, sphingolipids protected *Leishmania* from toxins and amphotericin. Thus, *Leishmania* organizes sterol-protective components differently from humans. Further, toxins and *Leishmania* serve as a system to understand fundamental rules governing sterol-protecting component membrane organization. We can use this information to help improve drugs targeting sterols.

## Introduction

Annually, 1.5-2 million cases and 70,000 deaths are caused by the neglected tropical disease, leishmaniasis. One first-line treatment for leishmaniasis is liposomal amphotericin B, which binds to membrane ergosterol and induces pores in the membrane of the causative parasites in the genus *Leishmania* (Baginski & Czub, 2009). However, the lipid environment enabling access of amphotericin B to the membrane is unknown. Resistance to amphotericin B is described for lab strains (Alpizar-Sosa *et al*, 2022), and amphotericin B has nephrotoxic side-effects (Berman, 2003; Coukell & Brogden, 1998), highlighting the need to understand the impact of lipid environment on amphotericin B and improve drug efficacy. One approach to improving amphotericin B efficacy could be to enhance its access to ergosterol and ability to damage the membrane. In order to achieve this goal, it is critical to understand the determinants that control sterol access.

Membrane sterol access is primarily controlled by sphingolipids in mammalian cells. In mammalian cells, plasma membrane cholesterol is evenly split between sphingomyelin-cholesterol complexes, other “inaccessible” cholesterol, and “accessible” cholesterol (Das *et al*, 2014). Accessible cholesterol is defined by the ability of the membrane cholesterol to bind to exogenous sterol binding agents, like cholesterol-dependent cytolysins (CDCs) (Das *et al*., 2014). Sphingomyelinases liberate cholesterol from cholesterol-sphingomyelin complexes, increasing sterol sensitivity to CDCs (Schoenauer *et al*, 2019). Thus, sphingolipids are a prominent factor governing the sterol accessibility in mammalian cells.

Compared to mammalian cells, *Leishmania* synthesize different lipids, which could alter sterol accessibility. Like cholesterol, ergosterol forms detergent resistant microdomains with the primary *Leishmania* sphingolipid, inositol phosphorylceramide (IPC) and GPI-anchored proteins like gp63 (Zhang *et al*, 2003). IPC comprises 10% of the total lipids in *Leishmania major* (Zhang & Beverley, 2010). Manipulation of IPC is best controlled at the level of its synthesis instead of cleavage. IPC is not cleaved by *B. cereus* sphingomyelinase, and the lipase that cleaves IPC (Zhang *et al*, 2009) likely cleaves other lipids, complicating the interpretation of any results using enzymes. Detection of IPC requires mass spectrometry approaches because it is not recognized by the animal sphingolipid sensors ostreolysin A or lysenin (Sepcic *et al*, 2004; Yamaji *et al*, 1998). Two key enzymes affecting IPC are serine palmitoyl transferase (SPT), the first committed step of sphingolipid synthesis, and IPC synthase (IPCS). Genetic ablation of the second subunit of SPT (*SPT2*) inactivates the enzyme. Knockout of *SPT2* prevents the formation of sphingosine, which leads to defects in ceramide and IPC synthesis and infectivity, but normal promastigote growth in logarithmic phase (Zhang *et al*., 2003). Similar results are observed when SPT2 is chemically inhibited with the drug myriocin (Zhang *et al*., 2003).

Similarly, genetic deletion of *IPCS* completely depletes IPC without impacting parasite growth or virulence (Kuhlmann *et al*, 2022). These mutants retain upstream sphingolipids, including enhanced ceramide levels (Kuhlmann *et al*., 2022). Thus, both genetic and pharmacological tools exist to measure the control of *Leishmania* sphingolipids over ergosterol accessibility.

Sterol accessibility is measured using inactive cholesterol-binding cytolysins (CDCs) from Gram positive bacteria. *S. pyogenes* secretes the CDC Streptolysin O (SLO), while *Clostridium perfringens* secretes the CDC perfringolysin O (PFO) (Thapa *et al*, 2020). These CDCs engage a range of sterols, including ergosterol (Savinov & Heuck, 2017). CDCs bind to sterol in the membrane, oligomerize and insert 20-30 nm pores into the membrane (Thapa *et al*., 2020). One advantage of CDCs is that their properties can be changed using mutagenesis, which we and others have extensively characterized (Farrand *et al*, 2015; Hotze *et al*, 2002; Johnson *et al*, 2012; Magassa *et al*, 2010; Mozola & Caparon, 2015; Ray *et al*, 2018; Romero *et al*, 2017).

Binding can be ablated by mutating two residues that drive sterol recognition (ΔCRM), or by introducing mutations that interfere with glycan binding (Mozola & Caparon, 2015). Similarly, mutation of two Gly to Val produce an inactive, non-toxic “monomer-locked” CDC that binds sterols (Hotze *et al*., 2002; Magassa *et al*., 2010; Ray *et al*., 2018). Finally, SLO and PFO engage sterols in distinct, but unknown, lipid environments (Farrand *et al*., 2015; Johnson *et al*., 2012). SLO binds to and inserts into membranes faster than PFO (Farrand *et al*., 2015; Johnson *et al*., 2012). The increased rate of binding and insertion is interpreted as SLO binding in a wider range of sterol microenvironments than PFO (Farrand *et al*., 2015; Johnson *et al*., 2012).

However, the identity of the microenvironment permissive for SLO or PFO remain unknown. The microenvironment requirements for each toxin can be switched to that of the other toxin by introducing point mutations in the membrane-binding L3 loop of the CDC (Farrand *et al*., 2015; Johnson *et al*., 2012; Ray *et al*., 2018). Thus, CDCs represent a versatile tool for probing the membrane environment and accessibility of sterols, such as the ergosterol environment in *Leishmania*.

Here, we tested the hypothesis that *Leishmania* shields the ergosterol target of amphotericin B in a manner similar to sterol shielding by mammalian cells. Surprisingly, we found that in contrast to mammalian cells, *Leishmania* sphingolipids did not prevent CDC binding, yet were able to prevent CDC cytotoxicity. Ceramide more strongly reduced PFO cytotoxicity, suggesting the mechanism of pore formation involves sensing ceramide via the L3 loop. Ceramide similarly contributed to the protection of *L. major* promastigotes from amphotericin B. Targeting ceramide and IPC may enhance ergosterol-targeting anti-leishmanial drugs.

## Results

### Sphingolipids do not limit accessible sterols in *L. major*

To determine if *L. major* sphingolipids shelter plasma membrane ergosterol, we measured the accessible ergosterol in wild type and sphingolipid-deficient *L. major* promastigotes. We used fluorescently labeled “monomer-locked” (ML) SLO and PFO, which detect accessible cholesterol in mammalian cells (Romero *et al*., 2017). Both SLO ML and PFO ML exhibited dose dependent binding to wild type (WT) *L. major* (Fig 1A, B), indicating they can bind to ergosterol in the *Leishmania* plasma membrane. Surprisingly, the sphingolipid-null *spt2^—^* (Zhang *et al*., 2003) showed no increase in SLO or PFO ML binding (Fig 1A, B). Overexpression of SPT2 in the *spt2^—^*background (*spt2^—^*/+SPT2) also did not change CDC binding (Fig 1A, B). These data suggest that sphingolipids do not shelter ergosterol in *L. major*.

**Figure 1.**
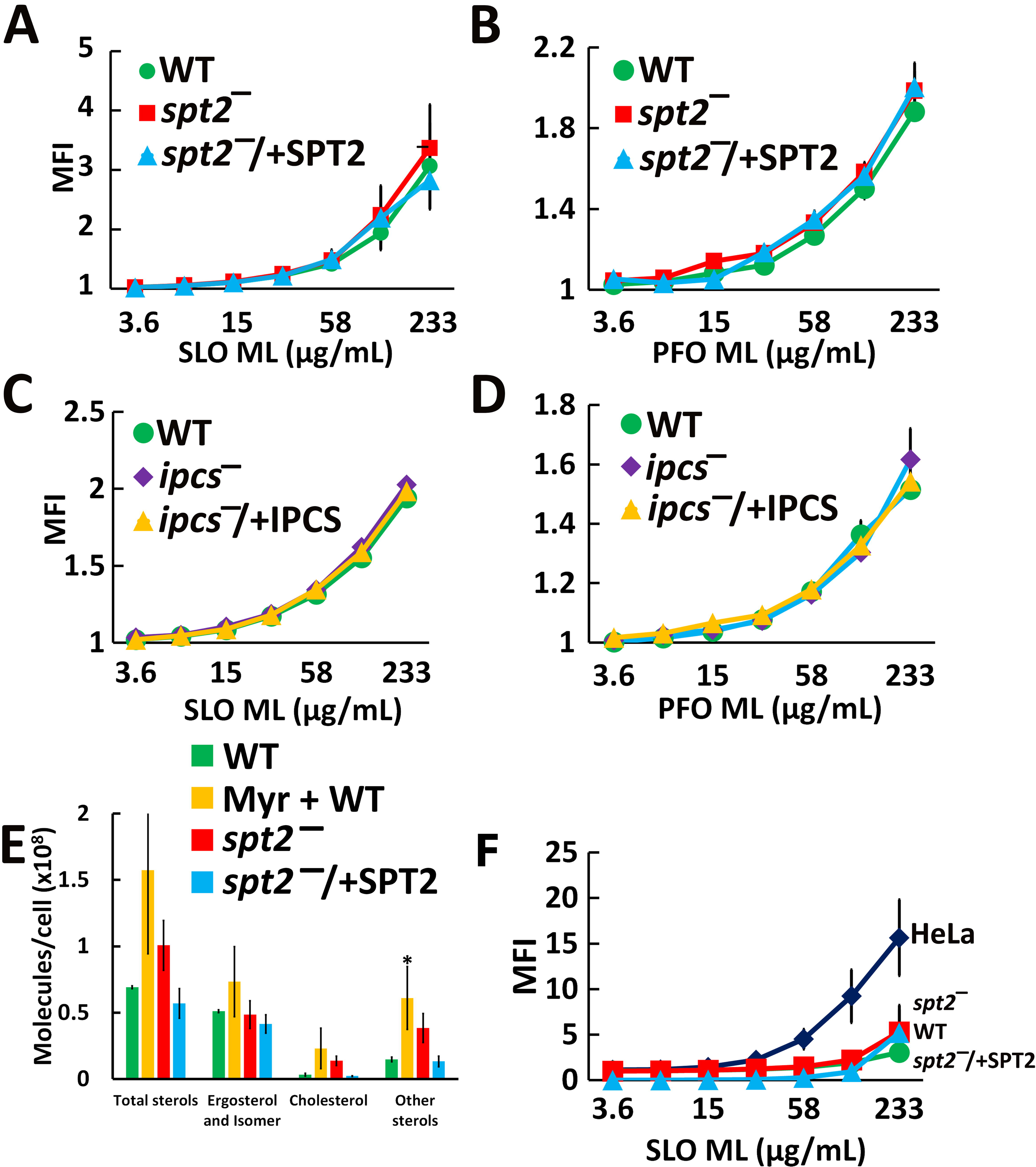
CDCs bind to *Leishmania major* promastigotes independently of sphingolipids. (A, B) Wild type (WT), *spt2^—^*, and *spt2^—^*/+SPT2, or (C, D) WT, *ipcs^—^*, and *ipcs^—^*/+IPCS *L. major* promastigotes were challenged with the indicated mass of (A, C) monomer-locked SLO (SLO ML) conjugated to Cy5 or (B, D) monomer-locked PFO (PFO ML) conjugated to Cy5 at 37°C for 30 min and analyzed by flow cytometry. (E) Total sterols from DMSO treated WT, *spt2^—^*, and *spt2^—^*/+SPT2, or WT treated with 10 µM myriocin *L. major* promastigotes were extracted, derivatized and analyzed by GC-MS. (F) HeLa cells or WT, *spt2^—^*, or *spt2^—^*/+SPT2 *L. major* promastigotes were challenged with SLO ML conjugated to Cy5 and analyzed by flow cytometry. Median Fluorescence Intensity x10 (MFI) of Cy5 fluorescence gated on live cells is shown. Graphs display the mean ± SEM of 3 independent experiments. * p < 0.05 by One-way ANOVA with Tukey’s multiple comparison post hoc testing (A-D, F) The x-axis is a log_2_ scale.

To confirm that sphingolipids do not shelter ergosterol in *L. major*, we used a *L. major* null mutant lacking the enzyme directly responsible for IPC synthesis from ceramide and phosphatidylinositol, *ipcs^—^*. Knockout of *IPCS* eliminates IPC, but retains ceramide (Kuhlmann *et al*., 2022). To complement the knockout, episomal expression of IPCS in *ipcs^—^* cells (*ipcs^—^* /+IPCS) was used (Kuhlmann *et al*., 2022). Similar to *spt2^—^*, SLO ML and PFO ML binding of *ipcs^—^* was equivalent to wild type and *ipcs^—^*/+IPCS *L. major* (Fig 1C, D). Prior characterization of the *ipcs^—^*cells revealed an absence of IPC and accumulation of ceramide in the *ipcs^—^* cells, which was reverted by complementation (Kuhlmann *et al*., 2022). We conclude that neither ceramide (absent in *spt2^—^*) nor IPC (absent in both) alter ergosterol accessibility to CDCs in *L. major*.

While the CDCs can access ergosterol independently of sphingolipids, the amount of sterol and the extent of accessibility remains unclear. It was previously reported that *spt2^—^ L. major* promastigotes have reduced ergosterol, but increased cholesterol levels (Armitage *et al*, 2018). To determine if interference in sphingolipid synthesis altered *L. major* sterol metabolism, we measured total sterols from promastigotes by gas chromatography-mass spectrometry (GC-MS). Promastigotes with inactivated *de novo* sphingolipid synthesis trended to an increase (30-100% more than WT control) in total sterols, but similar ergosterol and ergosterol isomer (like 5-dehydroepisterol) levels (Fig 1E). The sterol increase came from cholesterol (taken up from the media) and sterol biosynthetic intermediates (Fig 1E). To compare the extent of CDC binding between *L. major* and mammalian cells, we challenged both HeLa cells and *L. major* promastigotes with fluorescent SLO ML. Compared to HeLa cells, promastigotes poorly bound to SLO (Fig 1F). To determine if this was due to the smaller surface area of *L. major* promastigotes, we normalized MFI to surface area (Supplementary Fig S1A). When normalized to surface area, *L. major* bound more toxin per unit area than HeLa cells. Overall, these data suggest that CDCs sense ergosterol in *L. major* membrane independently of IPC or ceramide.

### *L. major* sphingolipids limit CDC cytotoxicity

Since CDCs bind poorly to *L. major* regardless of the sphingolipid composition, we tested if the CDC binding was sufficient to promote cytotoxicity. We challenged log phase wild type, *spt2^—^* or *spt2^—^*/+SPT2 promastigotes with SLO or PFO at 37°C for 30 min. We found that SLO killed wild type *L. major* only at very high doses, but SLO killed *spt2^—^* promastigotes at ∼28-fold lower dose (Fig 2A, Supplementary Fig S1B, C, S2A). Although PFO did not kill wild type or *spt2^—^*/+SPT2 promastigotes at any dose tested, it killed *spt2^—^*promastigotes like SLO (Fig 2B, Supplementary Fig S2B). We next tested the *ipcs^—^* and *ipcs^—^*/+IPCS, and found that *ipcs^—^* behaved similarly to *spt2^—^* (Fig 2C, D, Supplementary Fig S2C, D). We determined the CDC concentration needed to kill 50% of the cells (LC_50_) to compare the sensitivity of *spt2^—^*and *ipcs^—^ L. major*. To confirm our cytometry assay, we also performed an MTT assay. When killing was measured by MTT assay, we found similar results to our flow cytometry assay (Supplementary Fig S2E). This is consistent with our results in mammalian cells (Ray *et al*., 2018).

**Figure 2.**
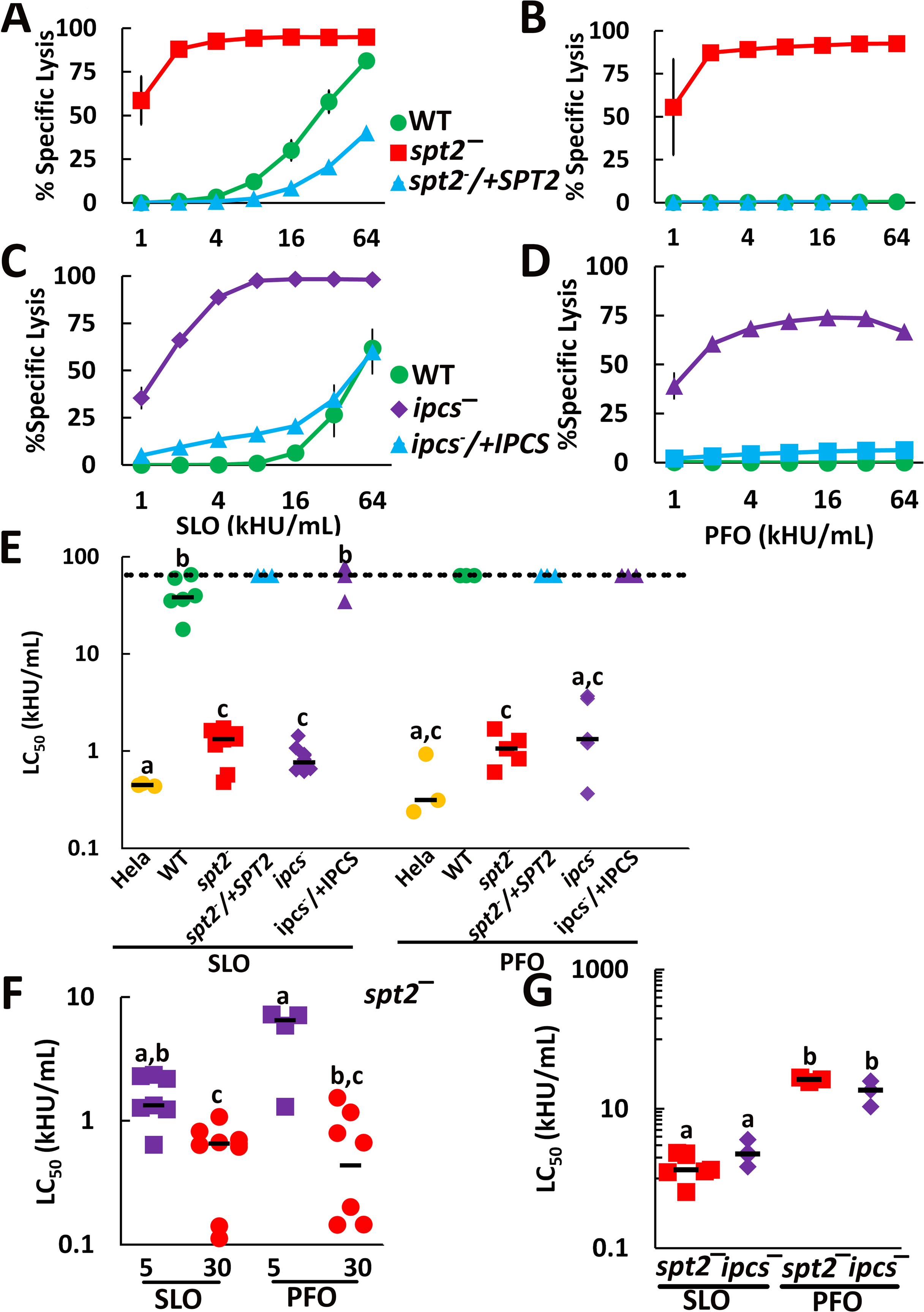
IPC protects *L. major* promastigotes from lysis by CDCs. (A, B) Wild type (WT), *spt2^—^*, and *spt2^—^*/+SPT2, or (C, D) WT, *ipcs^—^*, and *ipcs^—^*/+IPCS *L. major* promastigotes were challenged with (A, C) SLO or (B, D) PFO at the indicated concentrations for 30 min at 37°C and PI uptake measured by flow cytometry. (E-G) The LC_50_ was calculated as described in the methods after challenging the indicated genotypes of *L. major* with 31-4000 HU/mL or HeLa cells with 32-2000 HU/mL of SLO or PFO for 30 min (E, F) or with 62-4000 HU/mL SLO or 1000-64,000 HU/mL PFO for 5 min (F, G) and measuring PI uptake by flow cytometry. (A-D) Graphs display the mean ±SEM of at least 3 independent experiments. The dashed line indicates the highest concentration used. Points on this line had a LC_50_ value ≥64,000 HU/mL. (E-G) Graphs display individual data points and median from at least 3 independent experiments. Statistical significance was determined by one-way ANOVA with Tukey post-testing. Groups sharing the same letter were not statistically different. For example, HeLa cells challenged with SLO (group a) were not statistically distinct from HeLa cells challenged with PFO, or *ipcs^—^* cells challenged with PFO, but distinct from the remaining groups. In contrast, HeLa cells challenged with PFO (groups a and c), were only statistically distinct from WT and *ipcs^—^*/+IPCS L. major challenged with SLO (group b). (A-D) The x-axis is a log_2_ scale. (E-G) The y-axis is a log_10_ scale.

We next measured the LC_50_ for HeLa cells as a reference point because we have previously characterized cytotoxicity from CDCs in them (Keyel *et al*, 2011; Ray *et al*, 2022; Ray *et al*., 2018; Romero *et al*., 2017). Using HeLa cells as a reference point provides a relative estimate of how sensitive *L. major* promastigotes become when they are sphingolipid deficient. We found that sphingolipid-deficient *L. major* remained 1.5-2 times more resistant than HeLa cells when challenged with either SLO or PFO (Fig 2E, Supplementary Fig S2). Since sphingolipid-deficient *L. major* had elevated cholesterol levels, it is possible the sterol intermediates mediate the cytotoxicity. We challenged *L. major* promastigotes deficient in sterol methyltransferase (*smt^—^*) and C14 demethylase (*c14dm^—^*), which have normal sphingolipids, but elevated cholesterol levels (Mukherjee *et al*, 2020; Mukherjee *et al*, 2019), with SLO. We observed no killing of *smt^—^ L. major* promastigotes at doses ≤32,000 HU/mL, despite equivalent binding of toxin (Supplementary Fig S3A, B). Similarly, we observed no killing of *c14dm*^—^ *L. major* promastigotes at 4000 HU/mL (Supplementary Fig S3C). We conclude sterol alterations in sphingolipid-deficient *L. major* do not account for the observed cytotoxicity. Instead, sphingolipids protect *L. major* from CDC-mediated cytotoxicity.

One interpretation of the sphingolipid-mediated protection is that loss of sphingolipids destabilizes the membrane. While our previous results showing loss of sphingolipids did not perturb sterol-rich microdomains (Zhang *et al*., 2003), we tested this hypothesis using detergent challenge of sphingolipid sufficient and deficient cells. If the membrane is broadly destabilized, we predict the cells will be more sensitive to detergents like Triton X-100. When we challenged *L. major* promastigotes with Triton, we found that *c14dm^—^*, but neither *spt2^—^*, *c14dm^—^*/+C14DM, nor *spt2^—^*/+SPT2, were more sensitive to detergent than wild type promastigotes (Supplementary Fig S3D). Thus, we conclude that the sensitivity of sphingolipid-deficient promastigotes is specific to CDCs, and not the result of membrane destabilization.

To control for alternative mechanisms of membrane binding, any impurities in the CDC preparation, and the necessity of pore formation for *L. major* killing, we used three different non-hemolytic SLO mutants. The SLO ΔCRM lacks the cholesterol binding residues in domain 4 (Farrand *et al*, 2010). SLO Q476N cannot engage glycans on the cell surface needed for orientation (Mozola & Caparon, 2015; Shewell *et al*, 2020). These two mutants do not bind mammalian cells, but have no oligomerization or insertion defects (Farrand *et al*., 2010; Mozola & Caparon, 2015). If these toxins kill *spt2^—^* and *ipcs^—^ L. major*, it would indicate that SLO can engage *L. major* independently of glycans or sterol. If they fail to kill, it would indicate SLO needs these binding determinants to kill *L. major*. We used a third mutant toxin, SLO ML, which binds, but cannot form pores (Magassa *et al*., 2010; Romero *et al*., 2017). We challenged *spt2^—^* and *ipcs^—^ L. major* with SLO ΔCRM, SLO Q476N, SLO ML or wild type SLO (Supplementary Fig S2A, C, Supplementary Fig S3E, F). While SLO WT killed both *spt2^—^* and *ipcs^—^ L. major*, SLO ΔCRM, SLO Q476N, and SLO ML did not (Supplementary Fig S2A, C, Supplementary Fig S3E, F). Similarly, PFO ML failed to kill *spt2^—^* or *ipcs^—^* at all concentrations tested (Supplementary Fig S2B, D). These data indicate that CDCs require the same binding determinants to target both mammalian membranes and *L. major* membranes.

Since PFO and SLO showed different killing of wild type *L. major*, and have distinct membrane binding kinetics, we compared the rate at which SLO and PFO killed sphingolipid-deficient *L. major*. In mammalian cells, SLO more rapidly engages the membrane, whereas PFO binds more slowly (Farrand *et al*., 2015). Consequently, in HeLa cells there is a larger difference between PFO-mediated killing at 5 min and at 30 min than between SLO-mediated killing (Ray *et al*., 2018). We tested the sensitivity of *spt2^—^* or *ipcs^—^ L. major* after 5 min or 30 min of SLO or PFO challenge. For *spt2^—^*, we found that both CDCs had significant changes in LC_50_ between 5 min and 30 min (Fig 2F, Supplementary Fig S4A, B). While the SLO and PFO LC_50_ were similar at 30 min, at 5 min, there was a trend for less killing by PFO compared to SLO (Fig 2F). We next compared the *spt2^—^* and *ipcs^—^* mutants after CDC challenge. At 5 min, the *ipcs^—^*mutant was killed similarly to *spt2^—^*(Fig 2G, Supplementary Fig S4). These data indicate that sterol accessibility is insufficient to determine cytotoxic outcomes in *L. major*. Therefore, to sustain pore formation, SLO and PFO require different lipid environments in the *Leishmania* plasma membrane.

### Growth phase of promastigotes does not alter CDC sensitivity

The lipid environment fluctuations during the *L. major* growth cycle could impact CDC sensitivity. The *spt2^—^* promastigotes have defects in phosphatidylethanolamine (PE) synthesis when their growth reaches stationary phase (Zhang *et al*, 2007). In stationary phase, *L. major* promastigotes differentiate to the infectious metacyclic form over the course of approximately three days. We tested the impact of PE defects and metacyclogenesis on CDC killing of *spt2^—^ L. major*. We challenged *L. major* in log phase, or on day 1, 2 or 3 of stationary phase with 62-4000 HU/mL SLO or PFO. At the doses used, SLO and PFO killed *spt2^—^ L. major*, but did not kill wild type or *spt2^—^*/+SPT2 *L. major,* during stationary phase (Fig 3, Supplementary Fig S5A-F). The LC_50_ of SLO and PFO remained similar between log phase and throughout stationary phase (Figs 2, 3), suggesting that changes in PE synthesis did not impact CDC killing of *L. major*. We next determined if the differences we observed for the SLO LC_50_ at 5 min and 30 min in log phase were also present during stationary phase. The SLO LC_50_ for stationary phase promastigotes changed similarly to log phase (Fig 3B, C, Supplementary Fig S5G-I). Overall, the sensitivity of *spt2^—^*promastigotes to SLO and PFO did not significantly change during metacyclogenesis, suggesting that changes in PE during growth do not account for CDC sensitivity.

**Figure 3.**
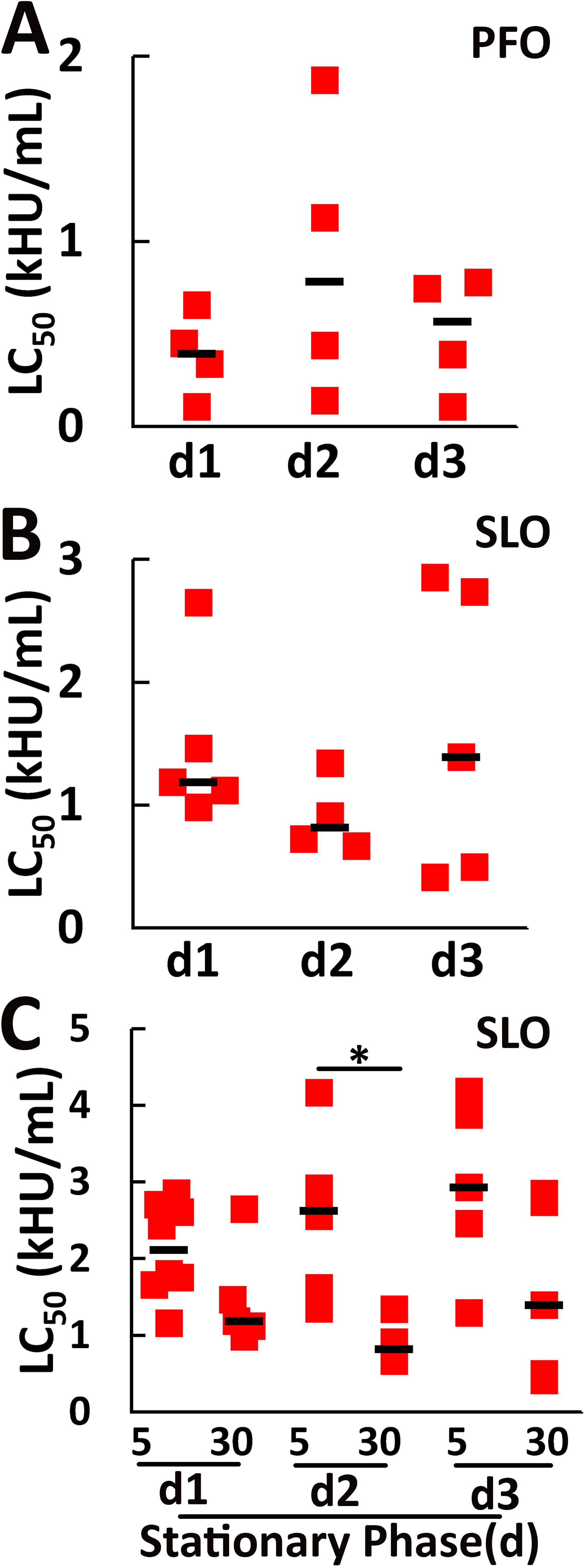
Transition to stationary phase does not alter CDC sensitivity of *spt2^—^ L. major* promastigotes. *spt2^—^ L. major* promastigotes in each day of stationary phase were challenged with (A) PFO, or (B, C) SLO for (A-C) 30 min or (C) 5 min at 37°C. PI uptake was analyzed by flow cytometry and LC_50_ calculated as described in the methods. Stationary phase d1, d2 and d3 represents 48, 72 and 96 hours post log phase. Graphs show individual data points and the median from at least 3 independent experiments. * p < 0.05 One way ANOVA was performed.

### Ceramide may hinder CDC cytotoxicity

One key difference between the *spt2^—^*and *ipcs^—^*that could account for the differences in cytotoxicity is ceramide. The *ipcs^—^*accumulates ceramide because it cannot synthesize it into IPC, whereas the *spt2^—^* cannot synthesize the precursors to ceramide. To determine the contribution of ceramide to CDC cytotoxicity, we used a chemical inhibitor of SPT, myriocin. We treated wild type, *spt2^—^*, *spt2^—^*/+SPT2, *ipcs^—^*, and *ipcs^—^*/+IPCS promastigotes with myriocin prior to CDC challenge. Myriocin increased the SLO sensitivity of wild type and *ipcs^—^*/+IPCS *L. major* similar to that of *spt2^—^*, confirming inhibitor efficacy (Fig 4, Supplementary Fig S6A-D).

**Figure 4.**
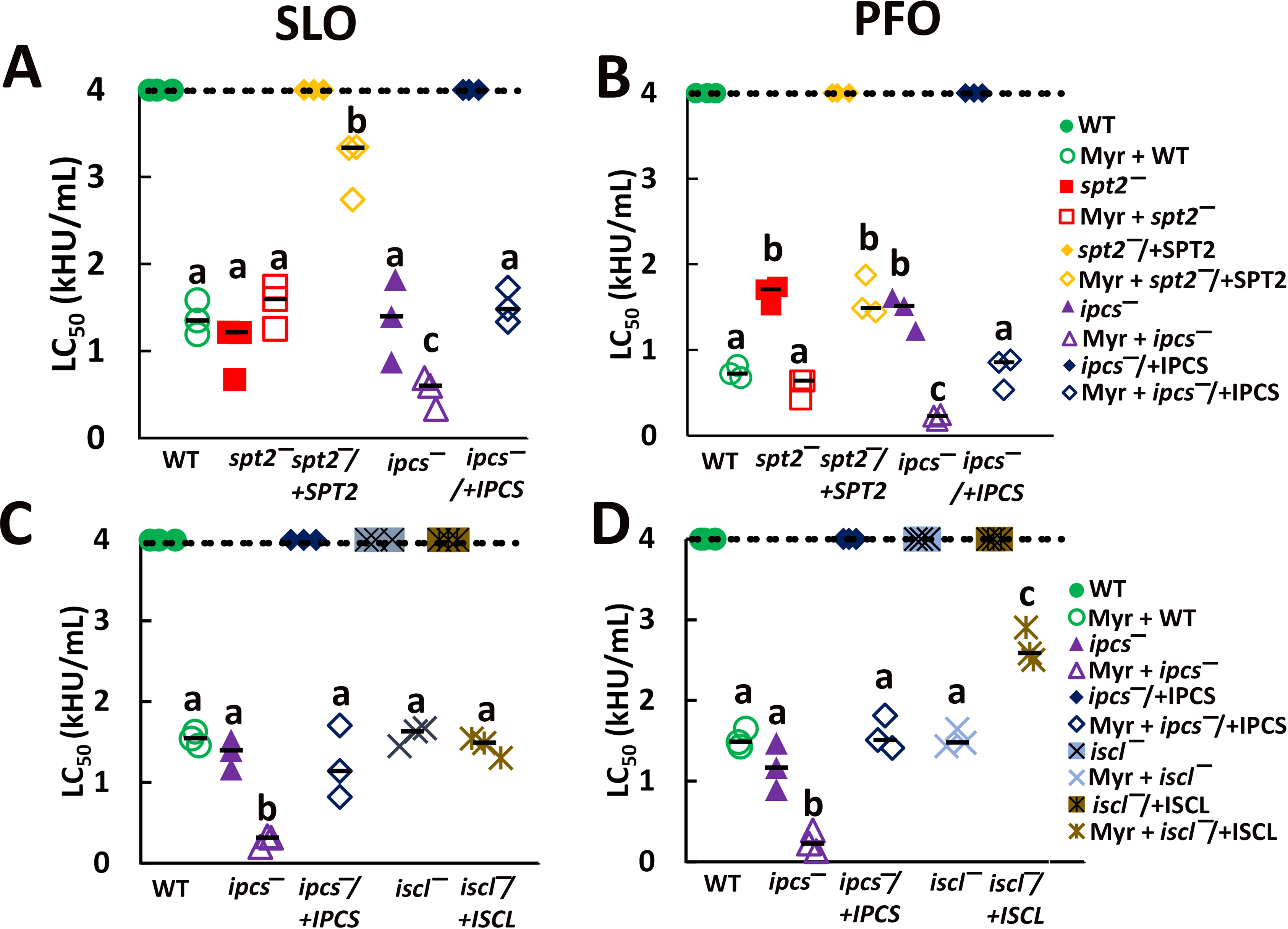
Myriocin treatment of *L. major* promastigotes suggests ceramide helps shield ergosterol. (A, B) Wild type (WT), *spt2^—^*, spt2*^—^*/+SPT2, *ipcs^—^* and *ipcs^—^*/+IPCS *L. major* promastigotes grown in either 10 μM Myriocin or DMSO supplemented M199 media were challenged with (A) SLO or (B) PFO for 30 min at 37°C. (C, D) WT, *iscl^—^*, *iscl^—^*/+ISCL*, ipcs^—^* and *ipcs^—^*/+IPCS *L. major* promastigotes grown in either 10 μM Myriocin or DMSO supplemented M199 media were challenged with (C) SLO or (D) PFO for 30 min at 37°C. PI uptake was analyzed by flow cytometry and LC_50_ calculated as described in the methods. WT, *ipcs^—^* and *ipcs^—^*/+IPCS represent distinct assays in each panel. Graphs display medians from 3 independent experiments. The dashed line indicates the highest concentration used. Points on this line had a LC_50_ value ≥4000 HU/mL. Statistical significance was determined by one-way ANOVA with Tukey post-testing. Groups sharing the same letter were not statistically different.

While sensitized to CDCs, myriocin-treated *spt2^—^*/+SPT2 were more resistant than myriocin-treated WT (Fig 4, Supplementary Fig S6). We attribute this resistance to elevated SPT2 levels in these promastigotes (Zhang *et al*., 2003). There was no change in the SLO LC_50_ for *spt2^—^* after myriocin treatment, indicating that myriocin was specific for SPT in our assay (Fig 4).

However, the LC_50_ for *ipcs^—^ L. major* decreased 2-fold for SLO and 6-fold for PFO upon myriocin treatment (Fig 4, Supplementary Fig S6). Furthermore, this increase in sensitivity surpassed that of either *spt2^—^*or *ipcs^—^*alone (Fig 4, Supplementary Fig S6). We interpret this finding to suggest that ceramide, which accumulates in *ipcs^—^*, but not *spt2^—^* promastigotes, contributes to protecting the *Leishmania* membrane from damage. Thus, ceramide and other perturbations in the lipid environment might contribute to protection from PFO.

Since PFO showed larger differences in *ipcs^—^ and spt2^—^* with and without myriocin, we interpret these findings to indicate that PFO is more sensitive to the overall lipid environment in the membrane than SLO. To test this interpretation, we challenged myriocin-treated, *I*nositol phospho*<u>S</u>*phingolipid phospholipase *<u>C</u>*-*<u>L</u>*ike (ISCL)-deficient (*iscl^—^*) *L. major* with CDCs. *iscl^—^ L. major* lack the lipase that converts IPC back into ceramide, and have elevated IPC levels (Zhang *et al*., 2009). In contrast to myriocin-treated *ipcs^—^ L. major*, which are expected to contain low levels of ceramide, myriocin-treated *iscl^—^ L. major* are expected to contain low levels of IPC. When challenged with SLO or PFO, myriocin-treated *iscl^—^* promastigotes phenocopied myriocin-treated WT promastigotes (Fig 4C, D, Supplementary Fig S6E-H). Consistent with the primary role of IPC in conferring resistance to cytotoxicity, untreated *iscl^—^* or *iscl^—^*/+ISCL *L. major* were resistant to SLO and PFO challenge, even at high toxin doses (Fig 4C, D, Supplementary Fig S6E-H). Notably, myriocin-treated *ipcs^—^* promastigotes were more sensitive to CDCs compared to WT or *iscl^—^* promastigotes (Fig 4C, D, Supplementary Fig SE-H). Overall, these data suggest that IPC provides the most protection against cytotoxicity, but ceramide may also protect *L. major* from the cytotoxicity of pore-forming toxins.

### The L3 loop in CDCs senses ceramide in the plasma membrane

We next determined the mechanism by which PFO is more sensitive to ceramide. One key difference between the membrane binding and accessibility of SLO and PFO is the amino acid sequence of the L3 loop (Farrand *et al*., 2015; Johnson *et al*., 2012). These differences in L3 loop can be switched by single point mutations. SLO S505D confers PFO-like binding and cytotoxicity on SLO, while PFO D434K has SLO-like binding and cytotoxicity (Farrand *et al*., 2015; Johnson *et al*., 2012; Ray *et al*., 2018). We tested if the cytotoxic differences we observed between CDCs— killing of wild type *L. major* at high toxin concentrations and the differences in *spt2^—^*and *ipcs^—^*LC_50_— could be explained by these L3 loop differences. We challenged wild type, *spt2^—^*, *ipcs^—^* and add-back *L. major* promastigotes with PFO, PFO D434K, SLO or SLO S505D for 30 min. We found that PFO D434K killed wild type and *spt2^—^*/+SPT2 *L. major* at high concentrations, similar to SLO, while PFO did not kill wild type *L. major* or the *spt2^—^*/+SPT2 (Fig 5A, B). SLO S505D phenocopied PFO, failing to kill wild type or *spt2^—^*/+SPT2 *L. major* (Fig 5A, B). While both PFO and PFO D434K killed *spt2^—^ L. major*, SLO S505D was less effective than either PFO or SLO in killing *spt2^—^ L. major* (Fig 5A, B). This is consistent with our previous findings in mammalian cells that SLO S505D has a higher LC_50_ than wild type SLO or PFO (Ray *et al*., 2018). We repeated these experiments using *ipcs^—^ L. major*. We found broadly similar results with *ipcs^—^* (Fig 5C, D). For each genotype, PFO D434K, with SLO-like binding, was the most potent toxin, and showed increased killing even in wild type *L. major* and *ipcs^—^*/+IPCS (Fig 5E). In contrast, SLO S505D, which has PFO-like binding, had decreased killing (Fig 5E). We conclude that the differences in SLO and PFO cytotoxicity are due to differential sensing of ceramide by the L3 loop.

**Figure 5.**
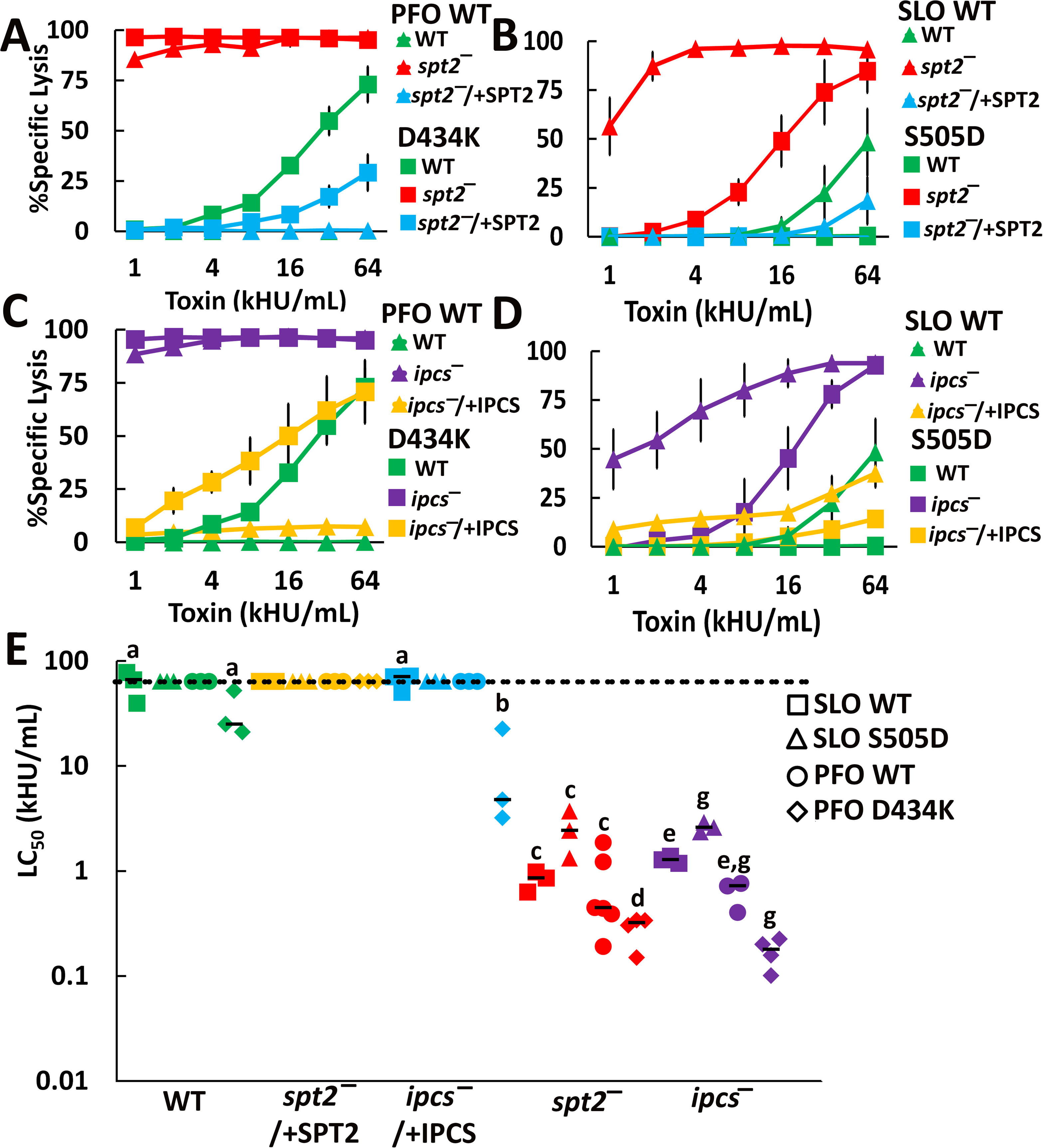
The CDC L3 loop controls CDC cytotoxicity against *L. major* promastigotes. (A, B) Wild type (WT), *spt2^—^*, and *spt2^—^*/+SPT2 or (C, D) WT, *ipcs^—^*, and *ipcs^—^*/+IPCS *Leishmania major* promastigotes were challenged with (A, C) PFO WT, PFO D434K, (B, D) SLO WT, or SLO S505D at the indicated concentrations for 30 min at 37°C. PI uptake was analyzed by flow cytometry and (E) LC_50_ calculated as described in the methods. (A-D) Graphs display the mean ±SEM of at least 3 independent experiments. (E) Graph displays individual data points and median from at least 3 independent experiments. The dashed line indicates the highest concentration used. Points on this line had a LC_50_ value ≥64,000 HU/mL. Both *spt2^—^* and *ipcs^—^* were statistically significant compared to WT or *ipcs^—^*/+IPCS by 2-way ANOVA. Statistical significance was determined by one-way ANOVA with Tukey post-testing. Within the same genotype, groups sharing the same letter were not statistically different. (A-D) The x-axis is a log_2_ scale. (E) The y-axis is a log_10_ scale.

### Ceramide and IPC protect *L. major* promastigotes from amphotericin B

Finally, these findings suggest that sphingolipid-deficient *L. major* promastigotes are more sensitive to amphotericin B. We first titrated both myriocin and amphotericin B against WT *L. major* promastigotes to determine the EC_50_. We found myriocin did not inhibit growth more than 25% promastigotes at 10 µM, while the EC_50_ of amphotericin B was 40 nM (Fig 6A, B). This is similar to the published EC_50_ of amphotericin B, which is 33 nM (Seifert & Croft, 2006). We next challenged log phase myriocin-treated WT and untreated WT, *spt2^–^*, *spt2^–^*/+*SPT2*, *ipcs^–^*, *ipcs^–^*/+*IPCS* promastigotes with amphotericin B. Removal of both IPC and ceramide increased the sensitivity of *L. major* 2-3-fold, whereas loss of only IPC was not sufficient to increase sensitivity to amphotericin B (Fig 6B-D). We conclude that both ceramide and IPC are needed to protect *L. major* promastigotes from amphotericin B.

**Figure 6.**
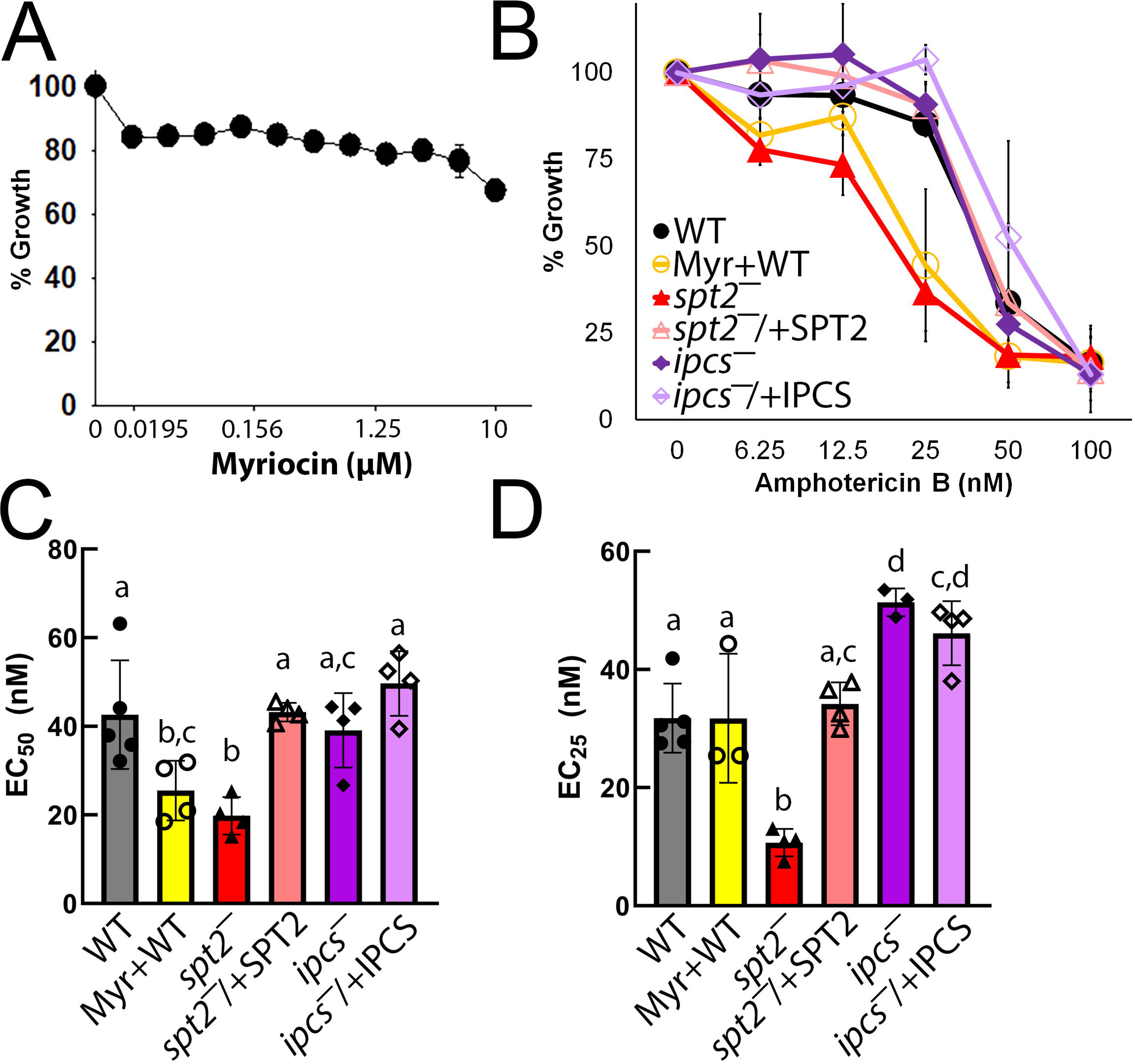
Ceramide protects *L. major* promastigotes from amphotericin B. (A) Wild type (WT) *L. major* promastigotes were challenged with the indicated concentrations of myriocin and growth determined relative to untreated. (B) Untreated WT, *spt2^—^*, and *spt2^—^*/+SPT2, *ipcs^—^*, *ipcs^—^*/+IPCS or 10 µM myriocin-treated WT *L. major* promastigotes were challenged with the indicated concentration of amphotericin B and growth determined relative to untreated WT. The (C) EC_50_ and (D) EC_25_ were calculated by logistic regression. Graphs display the mean ± SEM of 3 independent experiments. Statistical significance was determined by one-way ANOVA. Statistical significance (p<0.05) was assessed by post-hoc testing between groups. Statistical significance was determined by one-way ANOVA with Tukey post-testing. Groups sharing the same letter were not statistically different. (A, B) The x-axis is a log_2_ scale.

Based on these data, we propose a new model of CDC engagement with the *Leishmania* membrane. CDCs are able to bind to the membrane, independently of sphingolipids (Fig 7). In the absence of sphingolipids, both SLO and PFO oligomerize and insert into the membrane to kill the cell (Fig 7). The presence of ceramide without IPC provides limited protection against PFO, but not SLO. Recognition of ceramide in this environment is controlled by the L3 loop. Similarly, when Leishmania have their full complement of sphingolipids, ceramide precludes PFO toxicity at high doses, but not SLO (Fig 7). At lower toxin concentrations, however, the sphingolipids prevent toxin insertion by both CDCs.

**Figure 7.**
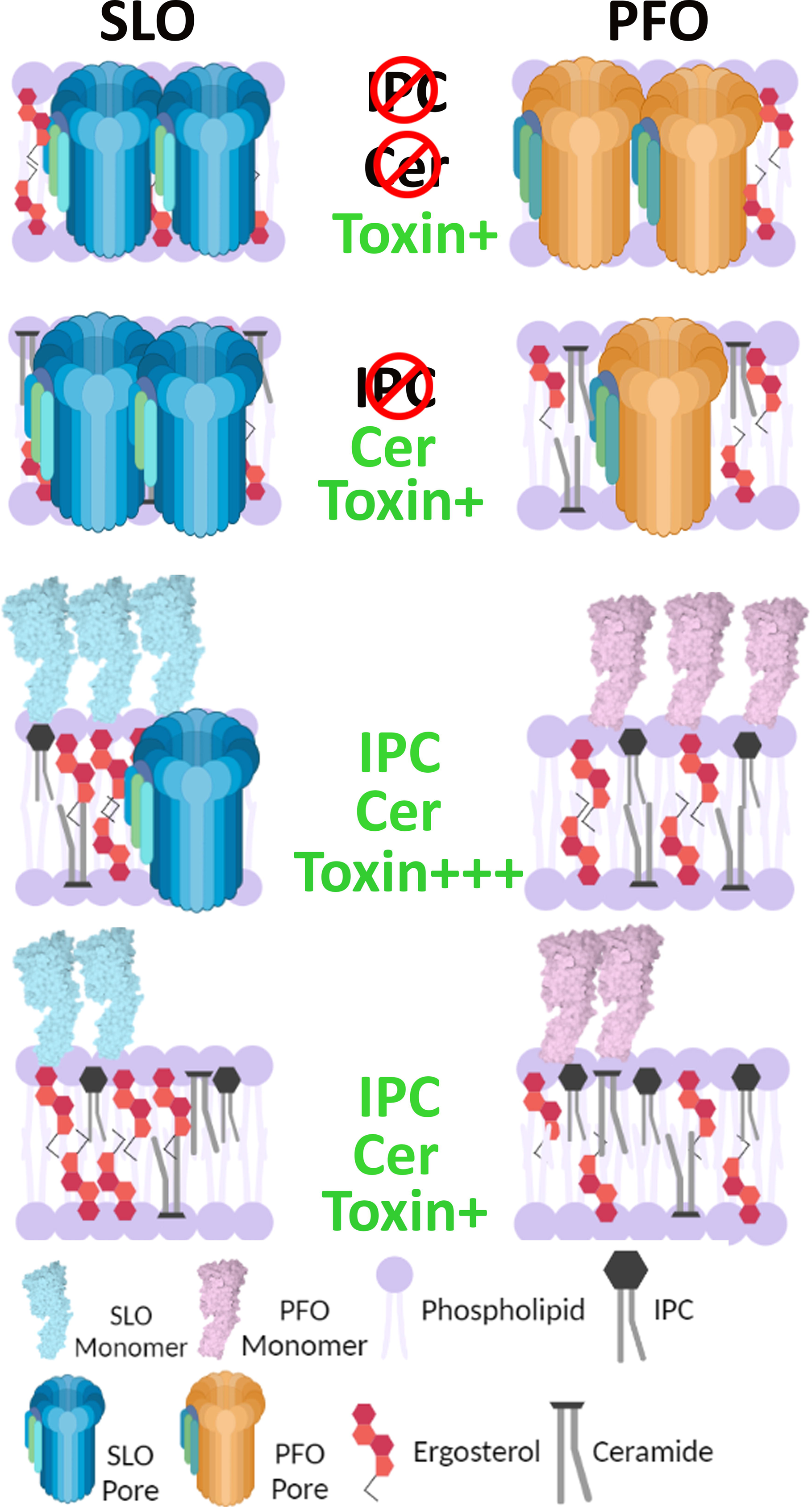
Cytotoxicity of CDCs towards *L. major* promastigotes is dependent on the presence of IPC. CDCs can bind to ergosterol in the membrane of *L. major* promastigotes regardless of sphingolipids. In the absence of IPC and ceramide, both SLO and PFO form lytic pores in the membrane. Elimination of IPC, but not ceramide, favors SLO pore formation, but reduced PFO pore formation. In the presence of ceramide and IPC, only SLO can still form pores at high doses (Toxin+++). At lower toxin doses (Toxin+), neither toxin can form pores in the membrane. The displayed inner-outer leaflet distributions of ergosterol and ceramide are for illustrative purposes only because the relative distribution of these lipids between inner and outer leaflet in *L. major* promastigotes is currently unknown.

## Discussion

Here we used the human pathogen *L. major* as a medically relevant model organism to understand toxin-membrane interactions. In contrast to mammalian systems, we found that *Leishmania* sphingolipids do not preclude CDC binding to sterols in the membrane, yet still interfere with cytotoxicity. Interference in toxicity was predominantly due to IPC. We further propose a mechanism where ceramide acts as one lipid to selectively reduce PFO, but not SLO, cytotoxicity in cells due to toxin binding determinants in the L3 loop. Disruption of both IPC and ceramide increased sensitivity of *L. major* to the anti-*Leishmania* drug amphotericin B. These data show that IPC and ceramide protect *L. major* from ergosterol-binding, pore-forming toxins. This study opens new horizons for future work on membrane repair in *Leishmania* and the competition between bacteria and *L. major*.

We found that Leishmania sphingolipids do not shelter ergosterol from CDCs. Ergosterol may not be sheltered from CDCs by IPC due to structural differences between sphingomyelin and IPC. Based on the crystal structure of the sphingomyelin/cholesterol-binding ostreolysin A and docking models with sphingomyelin, the choline headgroup flexibly lies across the membrane when sphingomyelin is complexed with cholesterol (Endapally *et al*, 2019). In contrast, the inositol headgroup of IPC is unlikely to adopt this conformation. This could account for the inability of IPC to prevent CDC binding to ergosterol.

We also considered the possibility that sphingolipid perturbations altered sterols, and altered sterols were responsible for the observed phenotypes. A previous study (Armitage *et al*., 2018) reported that *spt2^—^* promastigotes have lower ergosterol levels, and higher cholesterol levels. While we confirmed an increase in cholesterol and other sterol species with both genetic and chemical inhibition of SPT, we suggest this increase did not impact CDC binding or cytotoxicity. We observed no difference in CDC binding, suggesting that any sterol level alterations were insufficient to change CDC binding to the membrane. Since cholesterol is also involved in pore-formation (Giddings *et al*, 2003), we tested *L. major* promastigotes lacking SMT and C14DM (Mukherjee *et al*., 2020; Mukherjee *et al*., 2019). Both *smt^—^*and *c14dm^—^*have elevated total sterols, and accumulate cholesterol-type sterols, yet were not killed by SLO. Based on these considerations, we conclude that IPC and ceramide are primary determinants for CDC cytotoxicity, but not binding, to *Leishmania* membranes.

The failure to prevent binding by CDCs may represent different physiologic needs of *L. major* and mammalian cells. Mammalian cells need to sense sterol levels provided exogenously by the organism to modify sterol synthesis (Das *et al*., 2014), promote cellular signalling, and endocytosis (Rodal *et al*, 1999; Sen *et al*, 2011; Subramaniam & Johnson, 2002). In contrast, *Leishmania* promastigotes are free swimming organisms that synthesize their own sterols. Furthermore, no clear homologues of the sterol synthesis regulators sterol regulatory element-binding proteins (SREBP) or SREBP cleavage-activating protein (SCAP) have been detected in *Leishmania*. The lack of complications from SREBP signalling represents one advantage to using *Leishmania* as a model organism to understand sterol membrane dynamics.

While changes in *Leishmania* sphingolipid status did not interfere with CDC binding, changes in both IPC and ceramide reduced the ability of CDCs to kill *L. major*. This finding adds to the controversial role of sphingomyelin and ceramide in preventing vs accentuating damage in mammalian cells. On one hand, the sphingomyelinase/pore-forming toxin combination is evolutionarily conserved from bacteria like *C. perfringens* up to venoms in bees and snakes, suggesting that destroying sphingomyelin enhances toxicity. On the other hand, acid sphingomyelinase (Tam *et al*, 2010) and/or neutral sphingomyelinase (Schoenauer *et al*., 2019) have been reported to promote membrane repair. In mammals, it is not possible to separate the effects of sphingomyelin on cholesterol accessibility from the effects of sphingomyelin on directly interfering with CDC pore formation. Since IPC does not shelter sterol, we were able to determine that IPC interferes with pore formation after binding. We conclude that sphingolipids comprise one key component of the non-permissive membrane environment in *Leishmania* that reduces CDC pore-formation.

We used differences in SLO and PFO binding to mechanistically probe the non-permissive environment. SLO binds rapidly to membranes, whereas PFO requires more time (Farrand *et al*., 2015). This time difference is reflected in kinetic differences for PFO, but not SLO, cytotoxicity (Ray *et al*., 2018). Like mammalian cells, we observed kinetic differences in CDC cytotoxicity in *L. major*. Importantly, PFO was unable to find an environment to promote pore formation in *L. major*, even at supraphysiologic doses. This phenotype was reversed by point mutations in the L3 loop that switch the membrane specificity between PFO and SLO. Since PFO D434K regained the ability to kill wild type *L. major*, we conclude that this part of the L3 loop discriminates IPC and/or ceramide. Our findings support the idea that the L3 loop promotes membrane binding, and controls the lipid environment needed for pore formation (Farrand *et al*., 2015). If the L3 loop also discriminates sphingomyelin in mammalian cells remains to be determined.

Ceramide may modulate CDC cytotoxicity in *L. major*. We found significant differences between the extent of cytotoxicity in the *spt2^—^*and *ipcs^—^*, both of which lack IPC. We found that this difference was more pronounced when we blocked SPT with myriocin in the *ipcs^—^*, even though myriocin treatment phenocopied the *spt2^—^*in our system. The key difference in IPC blockade is that SPT is upstream of ceramide synthesis, whereas IPCS is downstream of ceramide. We suggest that IPC is more protective than ceramide, but ceramide also enhances protection. This contrasts with findings that addition of ceramide to liposomes containing 35% cholesterol increased membrane permeability to SLO (Zitzer *et al*, 2001). The differences may be due to the system used, protection via a ceramide-lipid/protein complex absent in liposomes, changes in membrane fluidity that permit CDC sensing of other lipids, or lipid packing or membrane architecture could be disrupted by sphingolipid or ceramide loss. One argument against these changes driving our observed phenotypes is previous work shows the proportion of detergent resistant membranes and global membrane architecture is unchanged in *spt2^—^ L. major* (Denny & Smith, 2004; Zhang *et al*., 2003). This suggests membrane architecture is broadly preserved. Second, binding did not change, which suggests loss of sphingolipids did not increase the extent of toxin binding. Overall, we propose that ceramide is one lipid species sensed by CDCs when forming pores in membranes.

While we showed that *Leishmania* sphingolipids limit cytotoxicity, but not binding, our study had limitations that can be explored in future studies. We did not determine the total fraction of plasma membrane ergosterol needed to support lysis by CDCs. In mammals, binding of inactive CDCs were used to probe the proportion of cholesterol in the membrane (Das *et al*., 2014), though the smaller size of *Leishmania* makes this more challenging to detect. Instead, we focused on comparing the accessible pool of sterol with the sphingolipid-sheltered pool of sterol in *Leishmania*. While we interpret our binding data to indicate sphingolipids do not protect ergosterol, an essential pool of ergosterol could exist that is bound to other proteins and/or lipids. We did not use a mouse model. The high efficacy of the single agents in the mouse model makes detecting synergistic effects from combination therapy there challenging.

We did not examine amastigotes because they salvage lipids from the host macrophage (Henriques *et al*, 2003; Moitra *et al*, 2021; Winter *et al*, 1994), the membranes are substantially similar in *spt2^—^*, *ipcs^—^*, and wild type amastigotes (Kuhlmann *et al*., 2022; Zhang *et al*, 2005), and contaminating macrophage membranes could obscure results. Future work is needed to determine how salvaged lipids impact sphingolipid-deficient *Leishmania* survival. Since CDC-producing Gram positive bacteria, including *S. pyogenes*, contribute to secondary bacterial infections during cutaneous leishmaniasis (Gimblet *et al*, 2017; Jayasena Kaluarachchi *et al*, 2021), CDCs could contribute to late stage infection, and/or the time required for resolution. Similarly, our work provides a foundation for understanding competition in the sandfly midgut between bacteria producing pore-forming toxins and *L. major* in the sandfly midgut.

Overall, we established *L. major* as a pathogenically relevant and genetically tractable model system for studying biological membranes, and identified key differences in sterol/sphingolipid organisation compared to mammalian cells. Most prior membrane biology work was done either in opisthokonts or in model liposomes, so there are few studies on biological membranes in other taxonomic groups, including human pathogens. We provide a blueprint for examining the membranes of non-standard organisms. These findings provide important perspectives for the generalization of biological membranes, especially compared to opisthokonts.

## Material and Methods

### Reagents

All reagents were from Thermofisher Scientific (Waltham, MA, USA), unless otherwise noted. Cysteine-less His-tagged PFO (PFO WT) in pET22 was a generous gift from Rodney Tweten (University of Oklahoma Health Sciences Center, Oklahoma City, OK, USA). Cysteine-less, codon-optimized SLO (SLO WT) in pBAD-gIII was synthesized at Genewiz (New Brunswick, NJ). The C530A mutant retains wildtype binding, hemolytic activity and pore structure, but it is redox resistant (Keyel *et al*, 2013). Monomer-locked (G398V/G399V) SLO, cysteine-less SLO (C530A), SLO S505D and PFO D434K were previously described (Ray *et al*., 2018; Romero *et al*., 2017). Glycan-binding (SLO Q476N) (Mozola & Caparon, 2015), and cholesterol binding (SLO T564A/L565A) (SLO ΔCRM) SLO mutants (Farrand *et al*., 2010), and monomer-locked (G298V/G299V) PFO mutants were generated using Quikchange PCR and verified by Sanger sequencing. Primer sequences are available upon request.

### Recombinant Toxins

Toxins were induced and purified as previously described (Keyel *et al*, 2012; Romero *et al*., 2017). Toxins were induced with 0.2% arabinose (SLO WT, SLO S505D, SLO G398V/G399V, SLO Q476N, SLO T564A/L565A (ΔCRM)), or 0.2 mM IPTG (PFO, PFO D434K, and PFO G298V/G299V) for 3 h at room temperature and purified using Nickel-NTA beads. For Cy5 conjugation, recombinant toxins were gel filtered into 100 mM sodium bicarbonate (pH 8.5) using a Zeba gel filtration column according to manufacturer’s instructions. Enough Cy5 monoreactive dye (GE Healthcare) to label 1 mg protein was added to 3-4 mg toxin and incubated overnight at 4°C. Conjugated toxins were gel filtered into PBS to remove unconjugated Cy5 dye, aliquoted, and snap-frozen in dry ice. Protein concentration was determined by Bradford assay and hemolytic activity was determined as previously described (Ray *et al*., 2018) using human red blood cells (Zen Bio, Research Triangle Park, NC, USA).

One hemolytic unit is defined as the amount of toxin required to lyse 50% of a 2% human red blood cell solution in 30 min at 37 °C in 2 mM CaCl_2_, 10 mM HEPES, pH 7.4, and 0.3% BSA in PBS. The specific activities of SLO monomer-locked Cy5 and PFO monomer-locked Cy5 were <10 HU/mg. They were used at a mass equivalent to wild-type SLO and PFO. Multiple toxin preparations were used (Table 1).

**Table 1.** Specific activity of active toxin preps used

| Toxin | Figure Used | Specific Activity (HU/mg) |
| --- | --- | --- |
| SLO WT | 2A, 2C, 2E, S2C | 1.23 x10<sup>6</sup> |
|  | 2E, S2A, S2F | 8.3 x10<sup>4</sup> |
|  | 2F, 2G, S4 | 1.42 x10<sup>5</sup> |
|  | 3B, S5D-F | 3.4 x10<sup>5</sup> |
|  | 3C, S5G-I | 1.11 x10<sup>5</sup> |
|  | 4, S6 | 3.3 x10<sup>5</sup> |
|  | 5 | 3.3 x10<sup>5</sup> |
|  | S1 | 4.92 x10<sup>5</sup> |
|  | S2E | 1.2 x10<sup>6</sup> |
|  | S3B | 3.77 x10<sup>5</sup> |
|  | S3C | 4.81 x10<sup>5</sup> |
|  | S3E-F | 2 x10<sup>5</sup> |
| SLO S505D | 5 | 3.95 x10<sup>4</sup> |
| PFO D434K | 5 | 7.87 x10<sup>6</sup> |
| PFO WT | 2B, 2D, S2D | 8.39 x10<sup>6</sup> |
|  | 2F, S2B, S2F | 2.73 x10<sup>4</sup> |
|  | 2F, S4 | 1.92 x10<sup>6</sup> |
|  | 2E | 1.63 x10<sup>5</sup> |
|  | 2G, S4, 4, S6 | 3.63 x 10<sup>5</sup> |
|  | 3, S5 | 4.57 x10<sup>6</sup> |
|  | 5 | 1.75 x10<sup>6</sup> |

### Leishmania strains and culture

LV39 clone 5 (Rho/SU/59/P) was used as the wild type strain, and all genetic mutants were made in this background. The serine palmitoyltransferase subunit 2 knockout Δspt2::HYG/Δspt2::PAC (*spt2^—^*), episomal addback Δspt2::HYG/Δspt2:PAC/+pXG-SPT2 (*spt2^—^*/+SPT2) (Zhang *et al*., 2003), the inositol phosphosphingolipid phospholipase C-like knockout Δiscl2::HYG/Δiscl2::PAC (*iscl^—^*), episomal addback Δiscl2::HYG/Δiscl2::PAC/+pXG-ISCL (*iscl^—^*/+ISCL) (Zhang *et al*, 2012), sterol methyltransferase knockout Δsmt::HYG/Δsmt::PAC (*smt^—^*) (Mukherjee *et al*., 2019), episomal addback Δsmt::HYG/Δsmt::PAC/+pXG-SMT (*smt^—^*/+SMT) (Mukherjee *et al*., 2019), sterol 14-α-demethylase knockout Δc14dm:HYG/Δc14dm::PAC (*c14dm^—^*) (Mukherjee *et al*., 2020), and episomal addback Δc14dm:HYG/Δc14dm::PAC/+pXG-C14DM (*c14dm^—^*/+C14DM) (Mukherjee *et al*., 2020) have been previously described. The inositol phosphorylceramide synthase knockout Δipcs::HYG/Δipcs::PAC (*ipcs^—^*) and episomal addback Δipcs::HYG/Δipcs::PAC/+pXG-IPCS (*ipcs^—^*/+IPCS) were generated in the Beverley lab (Kuhlmann *et al*., 2022). Elevation of ceramide levels and lack of IPC were confirmed by mass spectrometry (Kuhlmann *et al*., 2022). Wild type *L. major* LV39, *spt2^—^*, *ipcs^—^*, *smt^—^* and *iscl^—^* were cultured at 27°C in M199 medium with 0.182% NaHCO_3_, 40 mM Hepes, pH 7.4, 0.1 M adenine, 1 µg/mL biotin, 5 µg/mL hemin & 2 µg/mL biopterin and 10% heat inactivated fetal bovine serum (FBS), pH 7.4. Episomal addback cells *spt2^—^*/+SPT2, *ipcs^—^*/+IPCS, *smt^—^*/+SMT, and *iscl^—^*/+ISCL were maintained in complete medium in the presence of 10 µg/mL neomycin (G418) and 20 µg/mL blasticidin (BSD), except experimental passages.

Culture density and cell viability were determined by hemocytometer counting and flow cytometry after propidium iodide (PI) staining at a final concentration of 20 µg/mL. In this study, log phase promastigotes refer to replicative parasites at 2.0 – 8.0 x10^6^ cells/mL, and stationary phase promastigotes referred to non-replicative parasites at densities higher than 2.0 x10^7^ cells/mL. Cells are considered Stationary Day 0 when they reach 2.0 x10^7^ cells/mL. They are Stationary Day 1, Stationary Day 2 and Stationary Day 3 at 24, 48, 72 hours after they reach Stationary Day 0, respectively.

### Drug treatment of L. major

For myriocin treatment experiments, experimental log phase cells were seeded at 1.0 x10^5^ cells/mL in complete medium and either treated with 10 µM Myriocin dissolved in 1X DMSO (experimental) or an equivalent volume of diluent 1x DMSO (control). Cells were cultured and allowed to reach log phase in 48 hours before harvesting and processing cells for experiments. For amphotericin B, log phase *L. major* promastigotes were inoculated in complete M199 media at 2.0 × 10^5^ cells/ml in 0–100 nM of amphotericin B. Culture densities were measured after 48 hours of incubation in 24-well plates. EC_50_ and EC_25_ were determined by logistic regression using cells grown in the absence of amphotericin B as controls.

### Leishmania processing for experiments

Cells were cultured in complete medium to log phase or stationary phase, according to experimental requirements. Cells were counted, centrifuged at 3200 RPM (Rotor SX4750), 8 min to pellet cells at room temperature (25°C). Cells were washed with 1X PBS and counted again for accuracy. Cells were centrifuged at 3200 RPM (Rotor SX4750), 8 min to pellet cells at room temperature (25°C), and resuspended in serum free 1X M199 to a final concentration of 1.0 x10^6^ cells/mL. Thus, 100 µL of cells used per sample / well in 96 well plates contained 1.0 x10^5^ cells.

### HeLa cell culture

Hela cells (ATCC (Manassas, VA, USA) CCL-2) were maintained at 37°C, 5% CO_2_ in DMEM (Corning, Corning, NY, USA) supplemented with 10% Equafetal bovine serum (Atlas Biologicals, Fort Collins, CO, USA) and 1× L-glutamine (D10). They were negative for mycoplasma by microscopy.

### Binding assay with L. major promastigotes L. major

promastigotes were resuspended at 1 x10^6^ cells/mL in M199 media supplemented with 2 mM CaCl_2_ and 20 μg/mL propidium iodide. Cy5-conjugated toxins were diluted in serum free M199 media according to a mass equivalent to active toxin and further diluted in two-fold intervals. Cells were examined for PI and Cy5 fluorescence using an Attune flow cytometer. Debris was gated out and cells exhibiting high PI fluorescence (1-2 log shift) (PI high), low PI fluorescence (∼1 log shift) (PI low) or background PI fluorescence (PI neg) were quantified, normalized against untreated cells and graphed according to mass used for inactive toxin (Supplementary Fig S1B, C). Both PI neg and PI low populations remain metabolically active, indicating that only the PI high population are dead cells (Keyel *et al*., 2011). The median fluorescence intensity (MFI) of Cy5 labeled, PI negative population was quantified, background-subtracted using cells receiving no Cy5-conjugated toxin. MFI was plotted against mass of inactive Cy5-conjugated toxin (SLO ML or PFO ML). To normalize binding to surface area, we assumed HeLa cells in suspension were spheres of radius 7.5 µm, while *L. major* promastigotes were ellipsoids with major radius 5 µm and minor radii 1.25 µm. The ellipsoid surface area was calculated using the Knud-Thompson formula. The MFI was divided by the final calculated surface area and reported as normalized MFI (Supplementary Fig S1A).

### Flow cytometry cytotoxicity assay with Leishmania major promastigotes

Killing assays were performed as described (Haram *et al*, 2022). *L. major* promastigotes were resuspended at 1 x10^6^ cells/mL in M199 media supplemented with 2 mM CaCl_2_ and 20 μg/mL propidium iodide. HeLa cells were resuspended at 1x 10^6^ cells/mL in RPMI media supplemented with 2 mM CaCl_2_ and 20 μg/mL propidium iodide. Toxins were diluted in serum free M199 media for *Leishmania* promastigotes or in serum free RPMI media for Hela cells according to hemolytic activity (wild-type toxins) or equivalent mass (inactive mutant toxins) and further diluted in two-fold intervals. PI fluorescence in cells was measured using an Attune flow cytometer. Debris was gated out and cells exhibiting high PI fluorescence (1-2 log shift) (PI high), low PI fluorescence (∼1 log shift) (PI low) or background PI fluorescence (PI neg) were quantified, normalized against untreated cells and graphed according to toxin concentration (Supplementary Fig S1). Specific lysis was determined as follows: % Specific Lysis = (% PI High^Experimental^ - % PI High^Control^) / (100 - %PI High^Control^). The sublytic dose was defined as the highest toxin concentration that gave <20% specific lysis.

### MTT assay

The MTT assay was performed as described (Ray *et al*., 2018) with the following modifications. *L. major* promastigotes were resuspended at 2.5 x10^7^ cells/mL in phenol red-free DMEM supplemented with 2 mM CaCl_2_. Toxins were diluted in serum free, phenol red-free DMEM according to hemolytic activity and further diluted in two-fold intervals. In each well, 2.5 x10^6^ cells were incubated with toxin for 30 min at 37°C, washed, and incubated with 1.2 mM MTT reagent in phenol red-free DMEM for 4 h at 37°C. Formazan was solubilized overnight with SDS-HCl. Plates were read at A_450_. The % viability was determined as (A_450_xpt – background)/(A_450_control – background) x 100%. The specific lysis was calculated as 100 – % viability. The LC_50_ was determined from specific lysis curves using linear regression.

### Sterol analysis by gas chromatography/mass spectrophotometry (GC-MS)

Total lipids were extracted according to a modified Folch’s protocol (Folch *et al*, 1957). *L. major* promastigotes (DMSO treated or treated with 10 µM myriocin for 48 h first) were resuspended in chloroform: methanol (2∶1) at 1.0×10^8^ cells/mL along with the internal standard cholesta-3,5-diene [(FW = 368.84) from Avanti Polar Lipids] at 2.0×10^7^ molecules/cell and vortexed for 30 seconds. Cell debris was removed by centrifugation (2500 RPM/1000 g for 10 minutes) and supernatant was washed with 0.2 volume of 1X PBS. After centrifugation, the aqueous layer was removed and the organic phase was dried under a stream of N_2_ gas. Lipid samples were then dissolved in methanol at the equivalence of 1.0×10^9^ cells/mL. An internal standard, cholesta-3,5-diene (formula weight, 368.34), was provided at 2.0 × 10^7^ molecules/cell during extraction. For GC-MS, equal amounts of lipid extract were transferred to separate vial inserts, evaporated to dryness under nitrogen, and derivatized with 50 μL of N,O-Bis(trimethylsilyl)trifluoroacetamide plus 1% trimethylchlorosilane in acetonitrile (1:3), followed by heating at 70°C for 30 min. GC-MS analysis was conducted on an Agilent 7890A GC coupled with Agilent 5975C MSD in electron ionization mode. Derivatized samples (2 μL each) were injected with a 10:1 split into the GC column with the injector and transfer line temperatures set at 250 °C. The GC temperature started at 180°C and was held for 2 min, followed by 10°C/min increase until 300°C and then held for 15 min. To confirm that the unknown GC peak retention time matched that of the episterol standard, we also used a second temperature program started at 80°C for 2 min, ramped to 260 °C at 50°C/min, held for 15 min, and increased to 300°C at 10°C/min and held for 10 min. A 25-m Agilent J & W capillary column (DB-1; inner diameter, 0.25 mm; film thickness, 0.1 μm) was used for the separation.

### Phosphatidylethanolamine and sphingolipid analysis by Electrospray Mass Spectrometry

To analyze relative abundance of phosphatidylethanolamine and sphingolipids, parasite lipids were extracted using the Bligh–Dyer approach (Bligh & Dyer, 1959) and examined by electrospray mass spectrometry in the negative ion mode as previously described (Pawlowic *et al*, 2016).

### Statistics

Prism (Graphpad, San Diego, CA), Sigmaplot 11.0 (Systat Software Inc, San Jose, CA) or Excel were used for statistical analysis. Data are represented as mean ± SEM as indicated. The LC_50_ for toxins was calculated by linear regression of the linear portion of the death curve. Statistical significance was determined either by one-way ANOVA with Tukey post-testing, one-way ANOVA (Brown-Forsythe method) with Dunnett T3 post-testing, or Kruskal-Wallis, as appropriate. p < 0.05 was considered to be statistically significant. Graphs were generated in Excel and Photoshop (Adobe, San Jose, CA, USA).

## Acknowledgments

The authors would like to thank members of the Keyel and Zhang labs for critical review of the manuscript. We thank the College of Arts and Sciences Microscopy for use of facilities.

## Author Contributions

Conceptualization-CH, KZ, PAK; Formal Analysis-CH, PAK; Funding Acquisition-SMB, KZ, PAK; Investigation-CH, SM, RK, FMK, CF; Methodology-CH, SM, CF, FH, RK, PAK; Project Administration-KZ, PAK; Resources-FMK, FH, SMB, KZ, PAK; Supervision-PAK; Visualization-CH, PAK; Writing-Original Draft Preparation-CH, SM, RK, KZ, PAK; Writing-Review & Editing-CH, SM, RK, FMK, CF, FH, SMB, KZ, PAK

## Supplemental Figure Legends

**Supplementary Figure S1. Binding analysis and gating strategy for *Leishmania major* promastigotes.** (A) The MFI from Fig 1F was normalized by cell surface area, assuming a sphere of radius 7.5 µm for HeLa cells in suspension and an ellipsoid calculated using the Knud-Thompson formula with a = 5 µm and b = c = 1.25 µm. MFI was divided by surface area. (B) Total *Leishmania major* promastigotes are gated on R1 gate for SSC-H and FSC-H. *L. major* promastigotes are then gated for single cells (R2) using FSC-A and FSC-H. (C) From R2, *L. major* promastigotes are gated for fluorescence intensity of propidium iodide (PI) for dead cells and live cells. (A) The x-axis is a log_2_ scale. (C) Both axes are log_10_ scale.

**Supplementary Figure S2. Pore-formation, glycan- and sterol-binding determinants are all required for cytotoxicity in *Leishmania major* promastigotes.** (A, B) Wild type (WT), *spt2^—^*, and *spt2^—^*/+SPT2, or (C, D) WT, *ipcs^—^*, and *ipcs^—^*/+IPCS *L. major* promastigotes were challenged with (A) monomer-locked SLO (SLO ML), (B) monomer-locked PFO (PFO ML), (A, C) SLO or (B, D) PFO at the indicated concentrations for 30 min at 37°C and PI uptake measured by flow cytometry. (E) WT, *spt2^—^*, and *spt2^—^*/+SPT2 *L. major* promastigotes were challenged with 62-4000 HU/mL SLO for 30 min at 37°C and viability measured by MTT assay. Analysis is described in the methods. Points on the dotted line had an LC_50_ > 4000 HU/mL. (F) Hela cells were challenged with SLO WT or PFO WT at the indicated concentrations for 30 min at 37°C and PI uptake measured by flow cytometry. (A-D, F) The x-axis is a log_2_ scale.

**Supplementary Figure S3: Sterol alterations and general membrane perturbations do not account for cytotoxicity.** (A, B) WT, *smt^—^*, and *smt^—^*/+SMT, or *spt2^—^ L. major* promastigotes were challenged with (A) SLO ML conjugated to Cy5 or (B) SLO at the indicated concentrations for 30 min at 37°C and PI uptake measured by flow cytometry. (E) *spt2^—^* and (F) *ipcs^—^ L. major* promastigotes were challenged with SLO (WT), SLO Q476N or SLO ΔCRM at the indicated concentrations for 30 min at 37°C. (C, D) WT, *c14dm^—^*,*c14dm^—^*/+C14DM, *spt2^—^*, or *spt2^—^*/+SPT2 *L. major* promastigotes were challenged with (C) SLO or (D) Triton X-100 (%v/v) at indicated concentrations for 30 min at 37° C and PI uptake was analyzed by flow cytometry. Graphs display mean ± SEM of 3 independent experiments. **** p <0.0001 by one-way ANOVA compared to the other genotypes. The x-axis is a log_2_ scale.

**Supplementary Figure S4. Sphingolipid-deficient mutants of *L. major* are susceptible to CDCs.** (A, B) WT, *spt2^—^*, and *spt2^—^*/+SPT2, or (C, D) WT, *ipcs^—^*, and *ipcs^—^*/+IPCS *L. major* promastigotes were challenged with (A, C) SLO or (B, D) PFO at the indicated concentrations for 5 min at 37°C. PI uptake was measured by flow cytometry. Graphs display mean ± SEM of at least 3 independent experiments. The x-axis is a log_2_ scale.

**Supplementary Figure S5. Sensitivity of *spt2^—^ L. major* promastigotes to CDCs remains the same across three consecutive days of stationary phase.** Stationary phase wild type (WT), *spt2^—^* and *spt2^—^*/+SPT2 *L. major* promastigotes were challenged with (A-C) PFO for 30 min (D-F) SLO for 30 min or (G-I) SLO for 5 min at 37°C at indicated concentrations. PI uptake was analyzed by flow cytometry. Stationary phase (A, D, G) d1, (B, E, H) d2 and (C, F, I) d3 represents 48, 72 and 96 hours post log phase. Graphs display mean ± SEM of 3 independent experiments. The x-axis is a log_2_ scale.

**Supplementary Figure S6. Chemical blockade of SPT enhances CDC cytotoxicity against *L. major* promastigotes.** (A-D) Wild type (WT), *spt2^—^*, *spt2^—^*/+SPT2, *ipcs^—^*and *ipcs^—^*/+IPCS *L. major* promastigotes were grown in either 10 μM Myriocin or DMSO supplemented M199 media, and challenged with (A, C) SLO or (B, D) PFO for 30 min. PI uptake was measured by flow cytometry and specific lysis determined. (E-H) WT, *iscl^—^*, *iscl^—^*/+ISCL*, ipcs^—^* and *ipcs^—^*/+IPCS *L. major* promastigotes were grown in either 10 μM Myriocin or DMSO supplemented M199 media, and challenged with (E, G) SLO and (F, H) PFO for 30 min at 37°C. PI uptake was measured by flow cytometry and specific lysis determined. Graphs display mean ±SEM of 3 independent experiments. The x-axis is a log_2_ scale.

