## Supplemental Figures for "Sphingolipids protect ergosterol in the *Leishmania major* membrane from sterol-specific toxins"

### Supplementary Figure S1

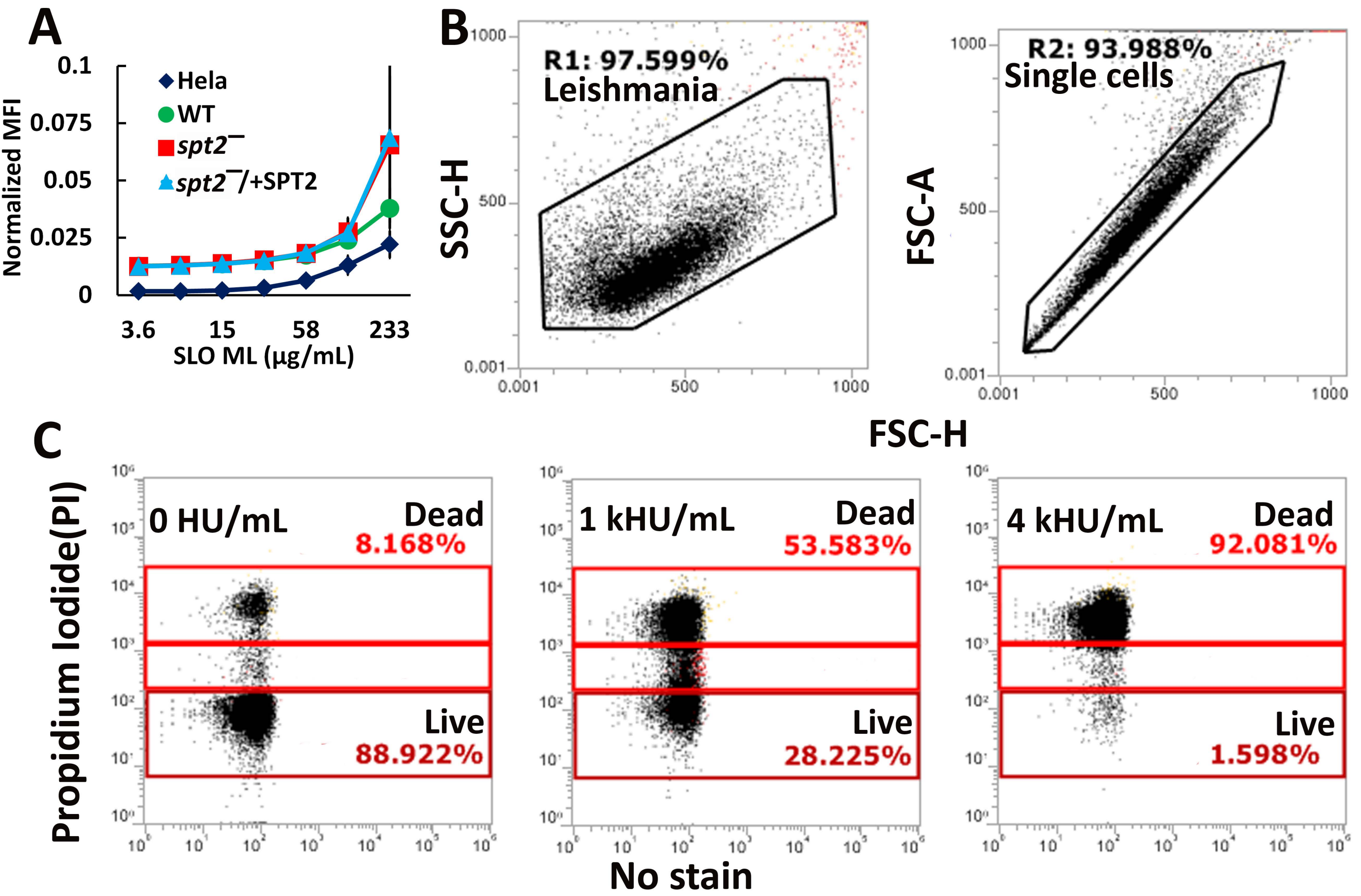

### Supplementary Figure S2

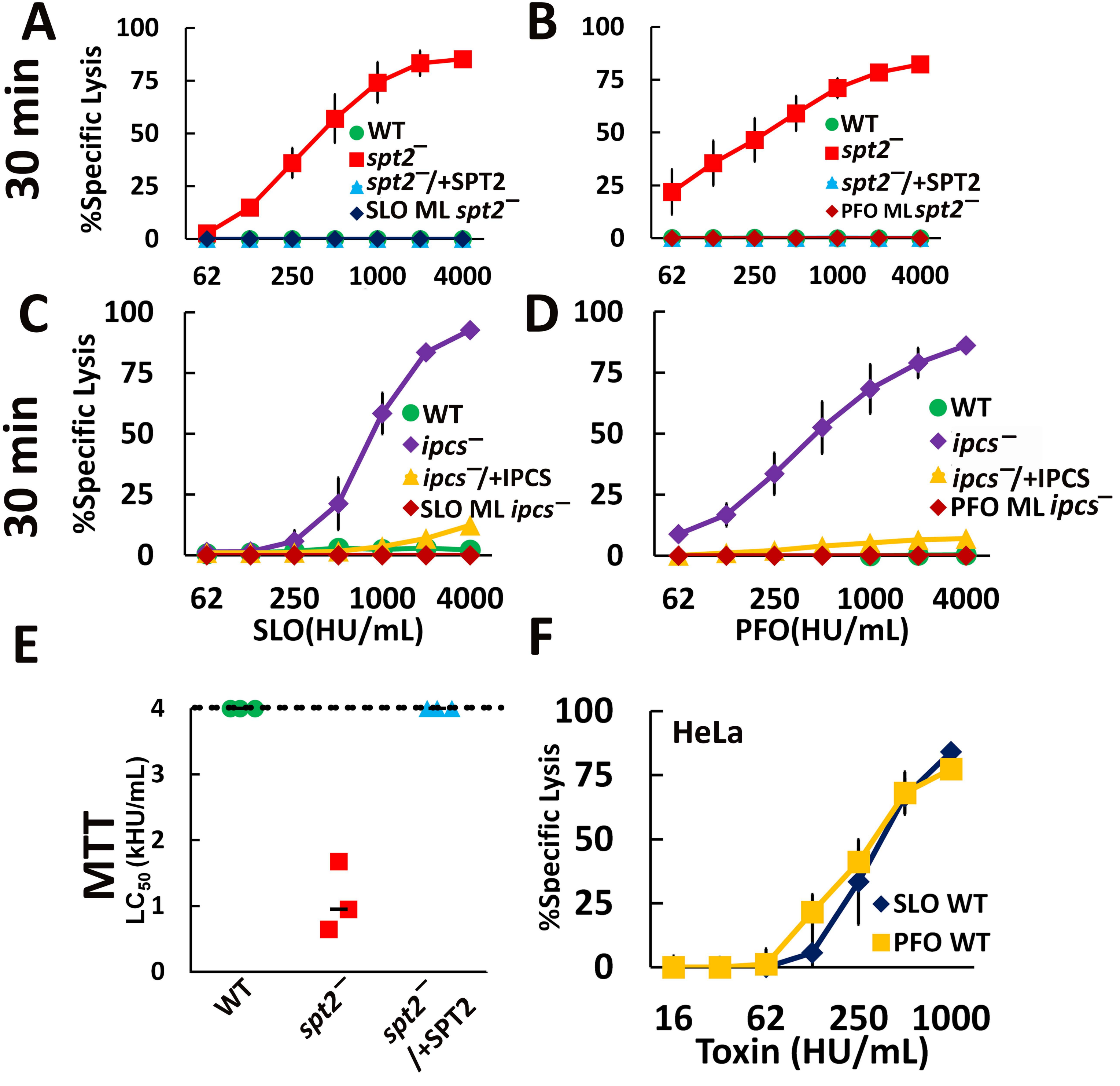

### Supplementary Figure S3

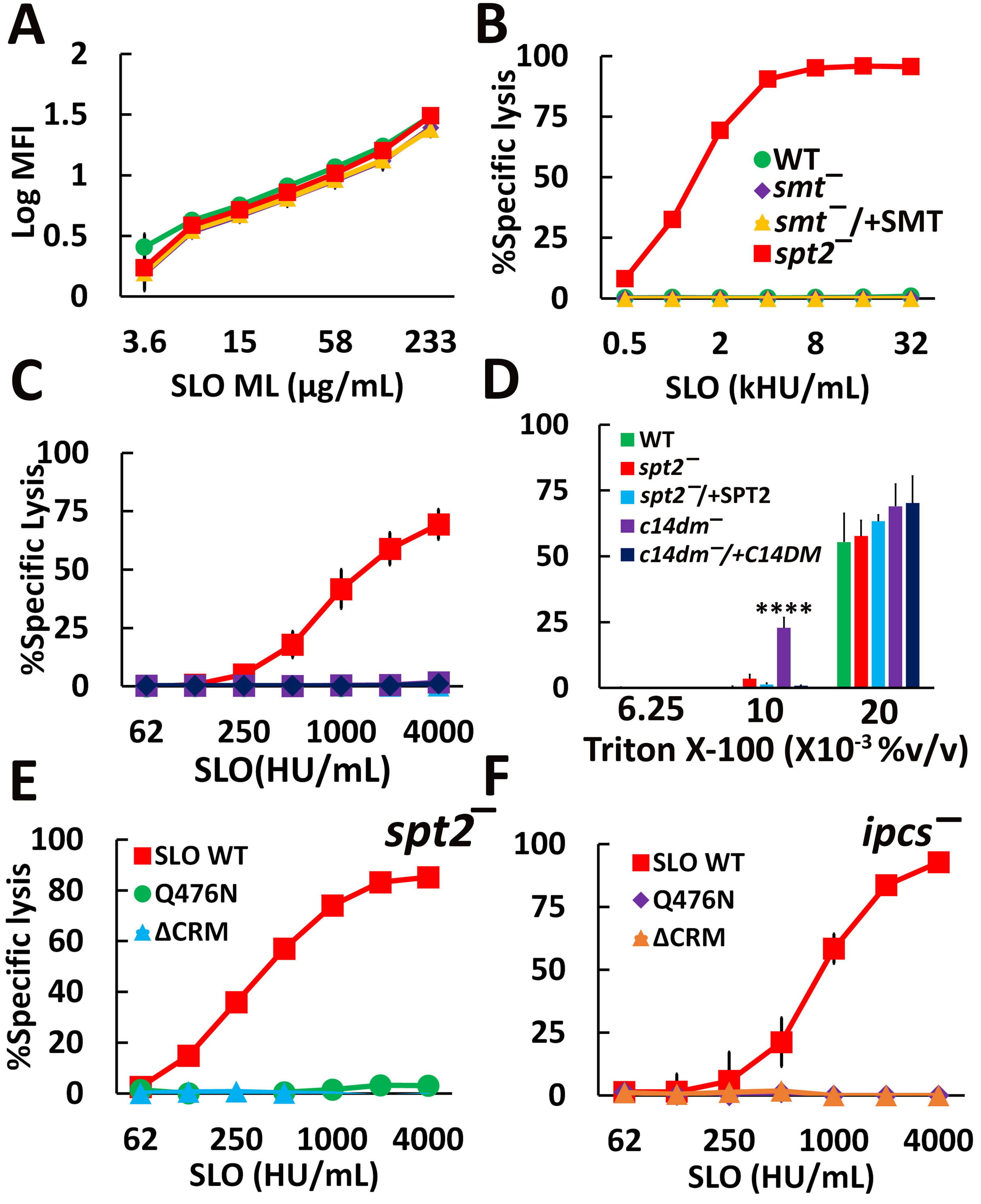

### Supplementary Figure S4

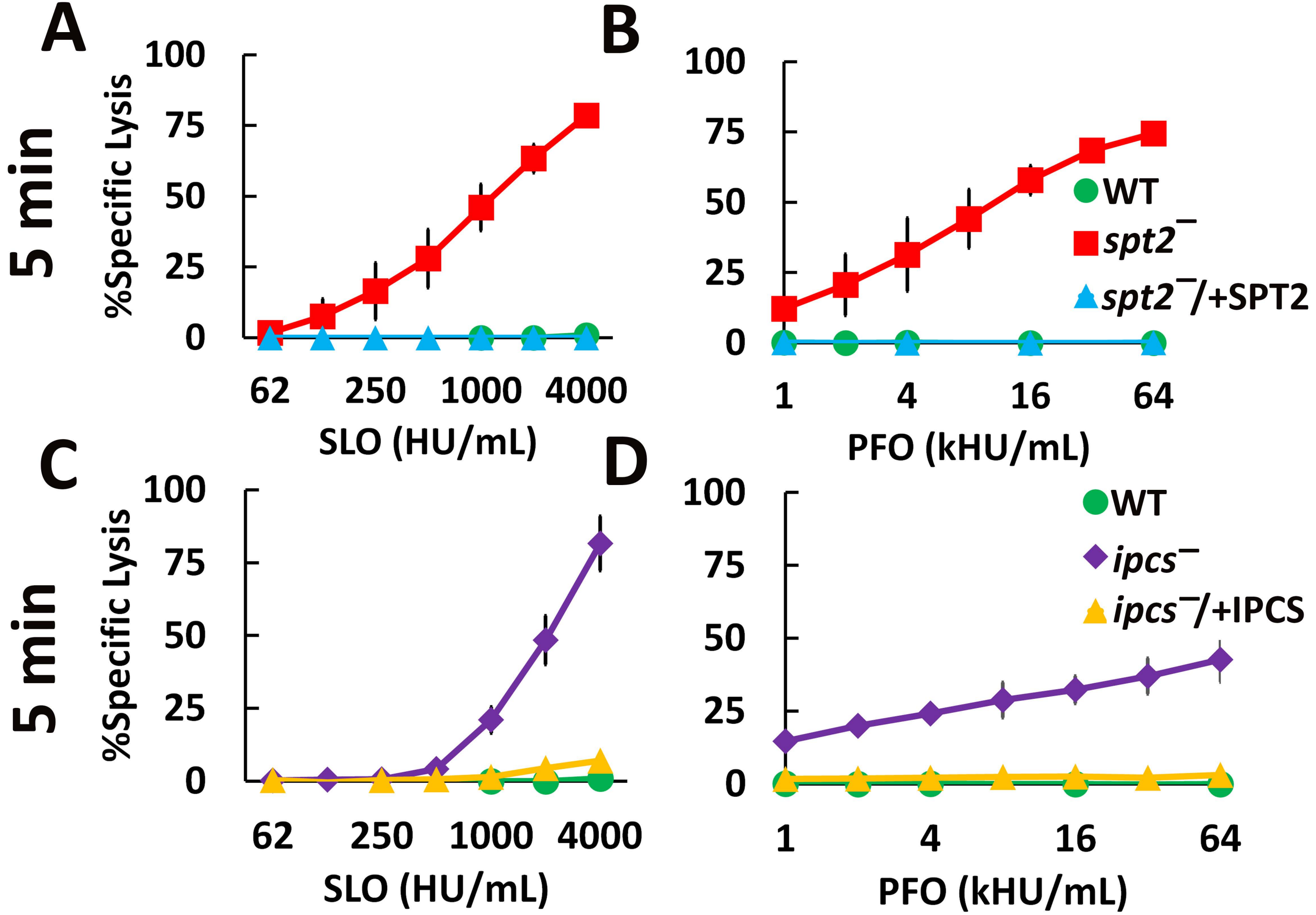

### Supplementary Figure S5

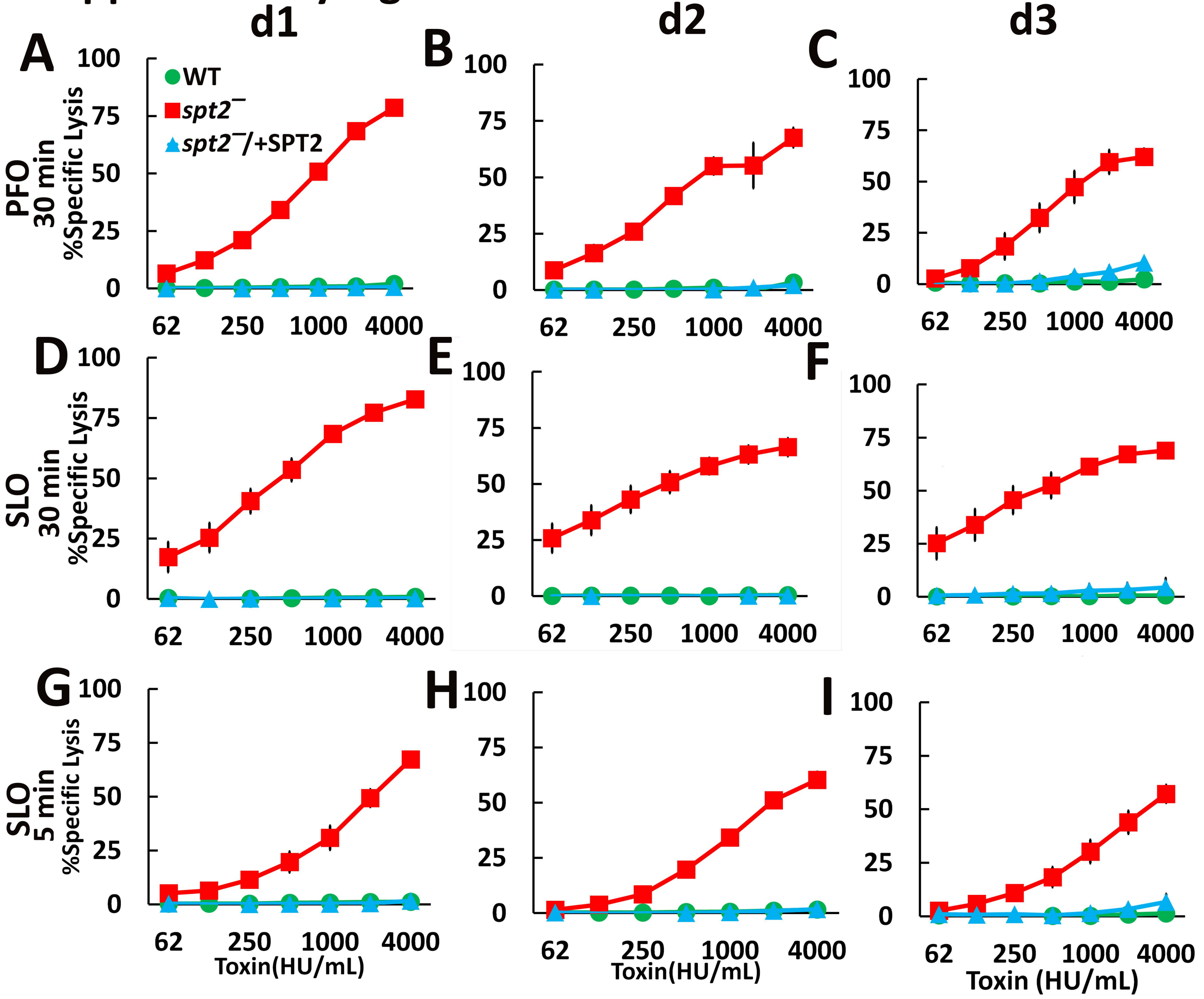

### Supplementary Figure S6

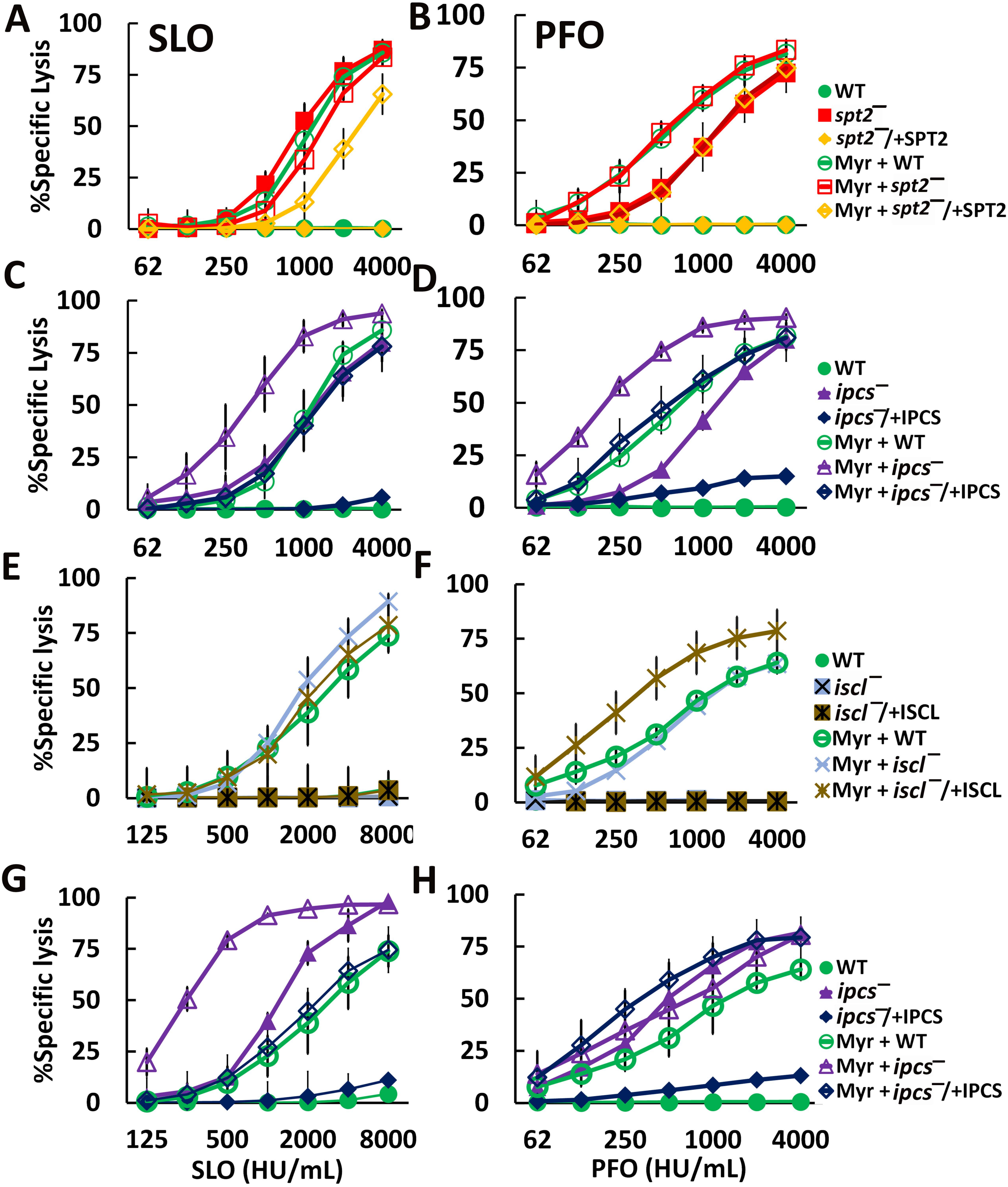
